# Impaired adult hippocampal neurogenesis in Alzheimer’s disease is mediated by microRNA-132 deficiency and can be restored by microRNA-132 replacement

**DOI:** 10.1101/2020.03.12.988709

**Authors:** Evgenia Salta, Hannah Walgrave, Sriram Balusu, Elke Vanden Eynden, Sarah Snoeck, Katleen Craessaerts, Nicky Thrupp, Leen Wolfs, Katrien Horré, Yannick Fourne, Alicja Ronisz, Edina Silajdžić, Zsuzsanna Callaerts-Vegh, Rudi D’Hooge, Dietmar Rudolf Thal, Henrik Zetterberg, Sandrine Thuret, Mark Fiers, Carlo Sala Frigerio, Bart De Strooper

## Abstract

**Summary:** Adult hippocampal neurogenesis (AHN) plays a crucial role in memory processes and is impeded in the brains of Alzheimer’s disease (AD) patients. However, the molecular mechanisms impacting AHN in AD brain are unknown. Here we identify miR-132, one of the most consistently downregulated microRNAs in AD, as a novel mediator of the AHN deficits in AD. The effects of miR-132 are cell-autonomous and its overexpression is proneurogenic in the adult neurogenic niche *in vivo* and in human neural stem cells *in vitro*. miR-132 knockdown in wild-type mice mimics neurogenic deficits in AD mouse brain. Restoring miR-132 levels in mouse models of AD significantly restores AHN and relevant memory deficits. Our findings provide mechanistic insight into the hitherto elusive functional significance of AHN in AD and designate miR-132 replacement as a novel therapeutic strategy to rejuvenate the AD brain and thereby alleviate aspects of memory decline.

## Introduction

Substantial evidence has demonstrated that neurogenesis occurs in adult human brain, is impaired in Alzheimer’s Disease (AD) patients and could contribute to the memory decline observed in AD pathology (Boldrini et al., 2018; Cipriani et al., 2018; Gatt et al., 2019; Kempermann et al., 2018; Lucassen et al., 2019, 2020; Moreno-Jiménez et al., 2019; Mu and Gage, 2011; Tartt et al., 2018; Tobin et al., 2019). Boosting adult hippocampal neurogenesis (AHN) was recently proposed as a putative therapeutic approach in AD (Choi and Tanzi, 2019; Choi et al., 2018; Lucassen et al., 2020), yet knowledge about the molecular mechanisms that can be used to leverage AHN is currently missing.

Cumulative data implicate adult neurons born at the subgranular zone of the dentate gyrus in a wide range of physiological mnemonic processes, including episodic, declarative, contextual and spatial memory, pattern separation, emotional control, functional forgetting and cognitive flexibility (Anacker and Hen, 2017; Baptista and Andrade, 2018; Toda et al., 2018). Pattern separation, a memory function that enables separation of similar representations into distinct, non-overlapping memories, is causally linked to the process of AHN and is functionally dependent on the adult-born neurons in the rodent dentate gyrus (Clelland et al., 2009; Danielson et al., 2016; Sahay et al., 2011). Several studies have reported impaired performance in pattern separation tasks in patients with amnestic mild cognitive impairment (MCI) with increased rate of progression to clinically probable AD (Petersen et al., 2001) and also in late-stage AD patients (Ally et al., 2013; Wesnes et al., 2014; Yassa et al., 2010). Interestingly, higher numbers of neuroblasts in the dentate gyrus correlate with better cognitive status in MCI patients (Tobin et al., 2019), while proliferation and differentiation capacity of adult neural stem cells have been correlated with memory performance in epilepsy patients (Coras et al., 2010). Taken together, these observations functionally link deficient AHN to memory decline in human neurological disorders and concomitantly put AHN forward as a promising therapeutic target in AD. However, a deeper understanding of the molecular regulators of AHN, and especially a better understanding of the molecular mechanisms going astray in disease, are a prerequisite for considering restoring AHN as a therapeutic strategy for neurodegenerative diseases. Previously, we and other have shown that miR-132 is strongly and reproducibly downregulated in the hippocampus of AD patients, notably early during disease progression (Hebert et al., 2013; Lau et al., 2013; Patrick et al., 2017; Pichler et al., 2017; Salta and De Strooper, 2017; Salta et al., 2016; Smith et al., 2015; Wong et al., 2013; Zhu et al., 2016). miR-132 overexpression in primary neurons or mouse brain represses pathological hallmarks of AD, such as amyloid plaques, TAU hyperphosphorylation and deposition, and neuronal death (El Fatimy et al., 2018; Hernandez-Rapp et al., 2015; Salta et al., 2016; Smith et al., 2015; Wang et al., 2017; Wong et al., 2013; Zhu et al., 2016) (reviewed in (Salta and De Strooper, 2017)). Whether the miR-132 deficiency in AD brain also contributes to pathophysiological events, such as reduced neurogenic potential or memory defects that precede late-stage protein aggregation and neuronal death, remains unknown.

Intriguingly, we have previously observed that miR-132 regulates the timing for cell cycle exit of radial glia-like neural stem cells in the developing vertebrate spinal cord (Salta et al., 2014), while miR-132 deletion in the dentate gyrus of young wild-type mice was shown to induce significant alterations in neurite maturation (Luikart et al., 2011; Magill et al., 2010). These data indicate a functional role of miR-132 in the neurogenic process, however, they do not provide further insights into disease mechanisms. Indeed, the question arises as to what extent loss of miR-132 expression is directly involved in the AHN defects observed in AD patients. Also, the more fundamental questions of whether and how miR-132 regulates AHN via direct or indirect effects on the neural stem cells of the adult niche, have not been addressed yet. Moreover, whether increasing miR-132 levels at the adult neurogenic niche would have beneficial effects on memory in AD in not known.

Here, we report that miR-132 is an integral and indispensable pro-neurogenic regulator of mouse and human neural stem cells, which operates cell autonomously via several pathways under physiological conditions and, importantly, also in AD, and that miR-132 replacement restores AHN and memory deficits in two distinct mouse models of AD by exerting a combination of beneficial effects in the adult hippocampal neurogenic niche.

## Results

### Adult hippocampal neurogenesis aberrations in AD parallel miR-132 deficiency in the dentate gyrus

In the subgranular zone, adult radial glia-like quiescent neural stem cells initially become activated, giving rise to transiently amplifying intermediate progenitors, which after limited rounds of proliferation generate fate-committed neuroblasts. Neuroblasts migrate tangentially along the subgranular zone and develop into immature neurons, which migrate radially into the granular cell layer to eventually mature into dentate granule neurons (Bond et al., 2015). Each of these activation and differentiation cellular stages can be labeled with distinct markers (Figure 1A).

**Figure 1.**
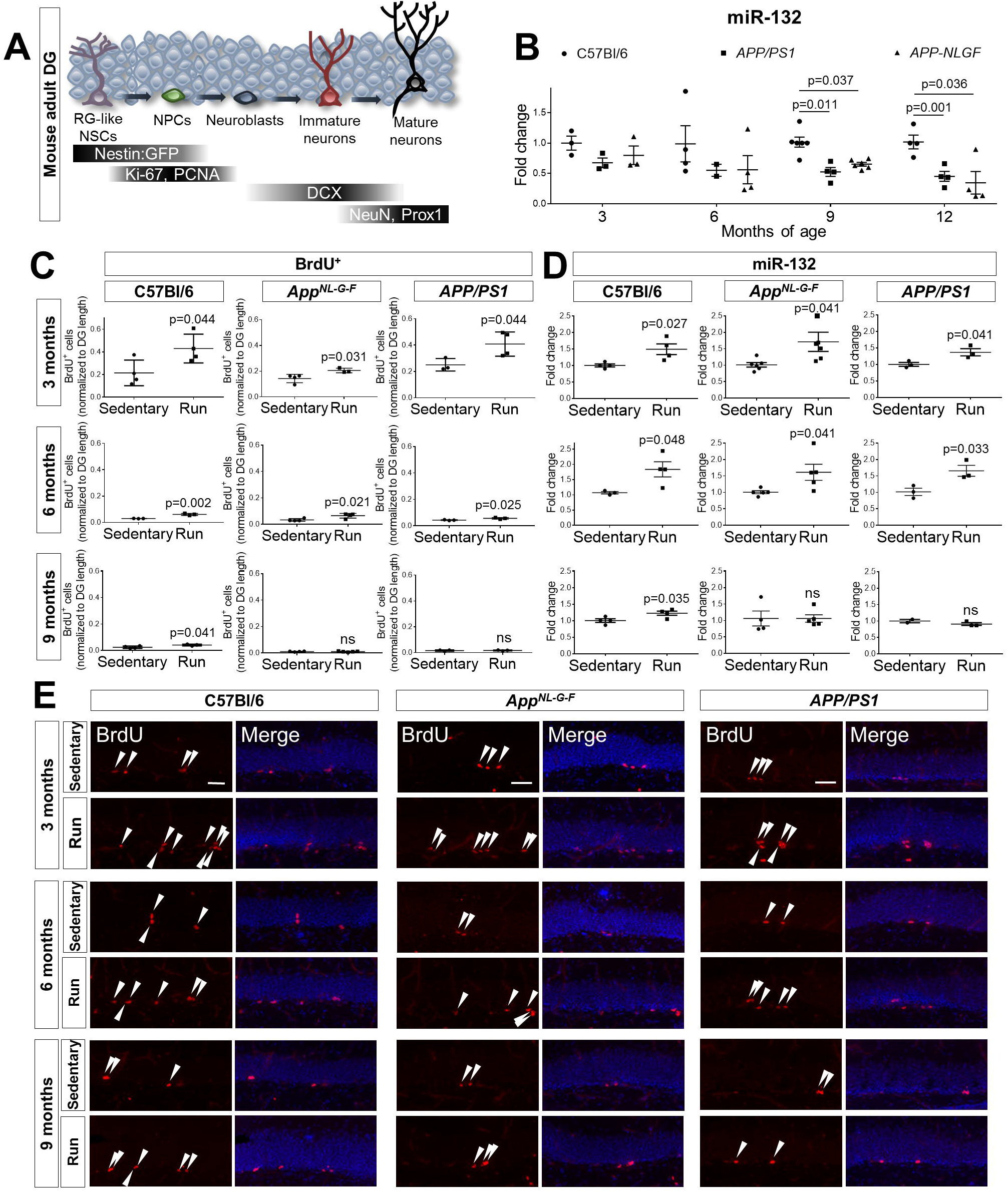
Adult neurogenesis is impaired by AD pathology and is associated with miR-132 levels. A. Schematic representation of the main fate transitions during adult hippocampal neurogenesis in the dentate gyrus (DG), including radial glia (RG)-like neural stem cells (NSCs), neuronal precursor cells (NPCs), neuroblasts, immature and mature neurons, as indicated by the expression pattern of distinct molecular markers. B. Semi-quantitative real-time PCR of miR-132 levels in the dentate gyrus of C57Bl/6, *APP/PS1* and *App^NL-G-F^* mice at 3, 6, 9 and 12 months of age. N=3-6 mice per time point. C. Quantification of experiment illustrated in panel E. BrdU^+^-cells in the subgranular zone of the dentate gyrus in C57Bl/6, *APP/PS1* and *App^NL-G-F^* control (sedentary) mice or mice exposed to voluntary running for 1 month (Run) at 3, 6 and 9 months of age. N=4-6 mice per group. D. Semi-quantitative real-time PCR of miR-132 levels in the dentate gyrus of C57Bl/6, *APP/PS1* and *App^NL-G-F^* sedentary control or running mice at 3, 6 and 9 months of age. N=4-6 mice per group. E. BrdU^+^-proliferating neuronal progenitors in C57Bl/6, *APP/PS1* and *App^NL-G-F^* sedentary control or running mice at 3, 6 and 9 months of age. Scale bars, 50 μm. Values are presented as mean ± SEM. In (B), two-way ANOVA with Tukey’s *post hoc* test for multiple comparisons were employed, while in (C, D), Student’s t-test was used. See also Figure S1 and Table S1.

To obtain an independent proof-of-concept of AHN deficiency in AD, we first performed a postmortem human brain immunohistochemical analysis of proliferating neuronal progenitors at the subgranular zone of the dentate gyrus in tissue sections from 10 non-demented control individuals and 10 AD patients (Figure S1A). Normalized proliferating cell nuclear antigen (PCNA)^+^-cell counts were significantly decreased by approximately 50% in AD dentate gyrus (Figure S1B). Astrocytes were negative for PCNA (Figures S1C, D), while PCNA^+^-microglia were excluded based on their larger and less spherical shape compared to PCNA^+^-neuronal precursor cells (Figures S1E, F). This reduction in the pool of proliferating cells, which eventually give rise to adult-born neurons possibly suggests an AHN-related deficit in the human AD brain. This observation is in line with recent reports providing strong evidence for decreased AHN in AD (Moreno-Jiménez et al., 2019; Tobin et al., 2019). Hence, miR-132 deficiency in the hippocampus of AD patients (Hebert et al., 2013; Lau et al., 2013; Patrick et al., 2017; Pichler et al., 2017; Salta and De Strooper, 2017; Salta et al., 2016; Smith et al., 2015; Wong et al., 2013; Zhu et al., 2016) is correlated with the decreased proliferation of neuronal precursors in the dentate gyrus.

We then asked whether miR-132 deficiency in AD is also functionally associated with the adult neurogenic niche. To address a possible genotype bias, we used two AD models, one overexpressing mutated forms of the human amyloid precursor protein (APP) and presenilin 1 (PS1) (*APP/PS1*) (Radde et al., 2006), and one carrying a knock-in mutated humanized *App* gene (*App^NL-G-F^*) (Saito et al., 2014). The dentate gyrus of male wild-type (C57Bl/6), *APP/PS1* and *App^NL-G-F^* mice was microdissected and miR-132 levels were measured at different ages by real-time PCR (Figure 1B). We observed an early trend for miR-132 downregulation in AD dentate gyrus compared to wild-type, which became significant at 9 months in both the *APP/PS1* and the *App^NL-G-F^* animals. Similar changes were observed in female mice (Figure S1G). Since synaptic loss in the hippocampus might be a confounding variable contributing to the reduction of synaptic miR-132 levels (Edbauer et al., 2010), we initially assessed the expression levels of syntaxin 1A (*Stx1A*) and synaptophysin (*Syp*) as a proxy for synaptic integrity by real-time PCR (Dickey et al., 2003). No changes in the levels of these transcripts were observed at 12 months of age in wild-type or AD mouse dentate gyrus (Figure S1H). To dissect more subtle synaptic alterations in the dentate gyrus of our AD mouse models, we analyzed co-localizing presynaptic (VGlut1-positive) and postsynaptic (PSD95-positive) puncta in the granular cell layer, the molecular cell layer and the hilar area at 3 and 9 months of age. Only a modest decrease of synaptic puncta was observed in 9-month old *APP/PS1* and *App^NL-G-F^* dentate gyrus compared to age-matched wild-type animals (Figures S1I-K), suggesting that the robust miR-132 decline in old AD dentate gyrus is not primarily induced by synaptic loss.

We next investigated whether AHN is affected in the two AD mouse models. We employed incorporation of a synthetic analog of thymidine, bromodeoxyuridine (BrdU), in the DNA of cells undergoing mitosis, in mice exposed to voluntary running, which is a widely used paradigm for induction of neuronal progenitor proliferation and adult neurogenesis (van Praag et al., 1999). One month of voluntary running increased the number of proliferating neuronal precursor cells at the subgranular zone of the dentate gyrus of both wild-type and AD mice compared to sedentary controls at 3 and 6 months of age (Figures 1C, E). However, the potential to induce neuronal progenitor proliferation at the subgranular zone upon running was compromised in 9-month old AD dentate gyrus, as opposed to wild-type mice (Figures 1C, E). Notably, this was paralleled by changes in miR-132 expression in the dentate gyrus. miR-132 levels significantly increased in running mice of all three genotypes at 3 and 6 months, yet no change was observed in running AD mice at 9 months of age (Figure 1D). Thus, adult neurogenesis and miR-132 induction drop in the aging AD mice. miR-212, a cognate microRNA of miR-132 which is co-transcribed from the same genetic locus, did not respond to physical exercise at any of the assessed time points in any of the genotypes (Figure S1L). Taken together, these results point towards a possible correlation between an amyloid plaque or amyloid beta (Aβ) oligomer pathology-induced effect on miR-132 expression and compromised AHN in two different mouse models for amyloid plaque pathology in Alzheimer’s disease.

### miR-132 expression at the adult hippocampal neurogenic niche is affected by amyloid pathology

We next assessed miR-132 localization in the adult neurogenic niche. One technical problem here is the strong baseline expression of miR-132 in the granule neurons of the dentate gyrus, which could mask the overall lower expression of miR-132 in cells at the subgranular zone (Luikart et al., 2011). We therefore used a nestin promoter-based green fluorescent protein (Nestin:GFP) reporter line in conjunction with additional markers to label neural stem cells and neuronal precursors at the subgranular zone of the dentate gyrus (Mignone et al., 2004) and employed fluorescence *in situ* hybridization and fluorescence-activated cell sorting (FACS) to determine the expression of miR-132. We use the term ‘Nestin:GFP^+^-niche cells’ to collectively refer to the GFP-positive dentate gyrus populations (Artegiani et al., 2017; Shin et al., 2015). Fluorescence *in situ* hybridization revealed that at basal conditions miR-132 is expressed in Nestin:GFP^+^/Ki-67^+^-proliferating neuronal precursor cells (Figure 2A) and Nestin:GFP^+^/Dcx^+^-immature neurons (Figure 2B) at the subgranular zone. Hybridization with a scrambled probe was used as a negative control (Figures S2A, B). Using a more quantitative method we next confirmed detection by real-time PCR and assessed the time course of miR-132 expression in Nestin:GFP^+^-niche cells isolated from wild-type mouse dentate gyrus. miR-132 levels were detectable in Nestin:GFP^−^- and to a smaller extent in Nestin:GFP^+^ -populations already at 1.5 month of age (Figure 2C). We observed a significant and specific increase of miR-132 expression in sorted cells at 12-months (Figures 2C and S2C). Interestingly, miR-132 upregulation did not occur in the Nestin:GFP^−^ -fraction which mainly consists of granule dentate neurons (Figure 2C), indicating that miR-132 is upregulated in the neurogenic niche, but not in the mature neurons, over time. Moreover, induction of adult neurogenesis via physical exercise elicited a significant upregulation of miR-132 only in the Nestin:GFP^+^-niche cell fraction (Figures 2D) and not in the Nestin:GFP^−^-neuronal population (Figure 2E). This upregulation was specific for miR-132 as the levels of miR-212 and of miR-124, a brain-enriched microRNA, were not changed (Figures S2D-G). We then asked whether amyloid plaque pathology affects miR-132 levels at the hippocampal neurogenic niche. To address this question, we crossed Nestin:GFP with *App^NL-G-F^* (Nestin-NLGF) mice and measured miR-132 levels in GFP^+^- and GFP^−^-cells isolated from the dentate gyrus at different ages. miR-132 levels in the Nestin:GFP^+^-niche cells of the AD mouse dentate gyrus increased at 6 months of age, similarly to wild type mice, but sharply decreased at 9 months (Figures 2F, S2I) coinciding with the inability of the neuronal progenitors in the AD mouse model brain to proliferate upon running. No significant changes were observed for miR-212 (Figure S2H). Taken together, these data show that miR-132 is recruited by adult neural stem cells and progenitors as part of the response to exercise- or aging-related stimuli, but fails to do so upon exposure to Aβ pathology in the hippocampal neurogenic niche.

**Figure 2.**
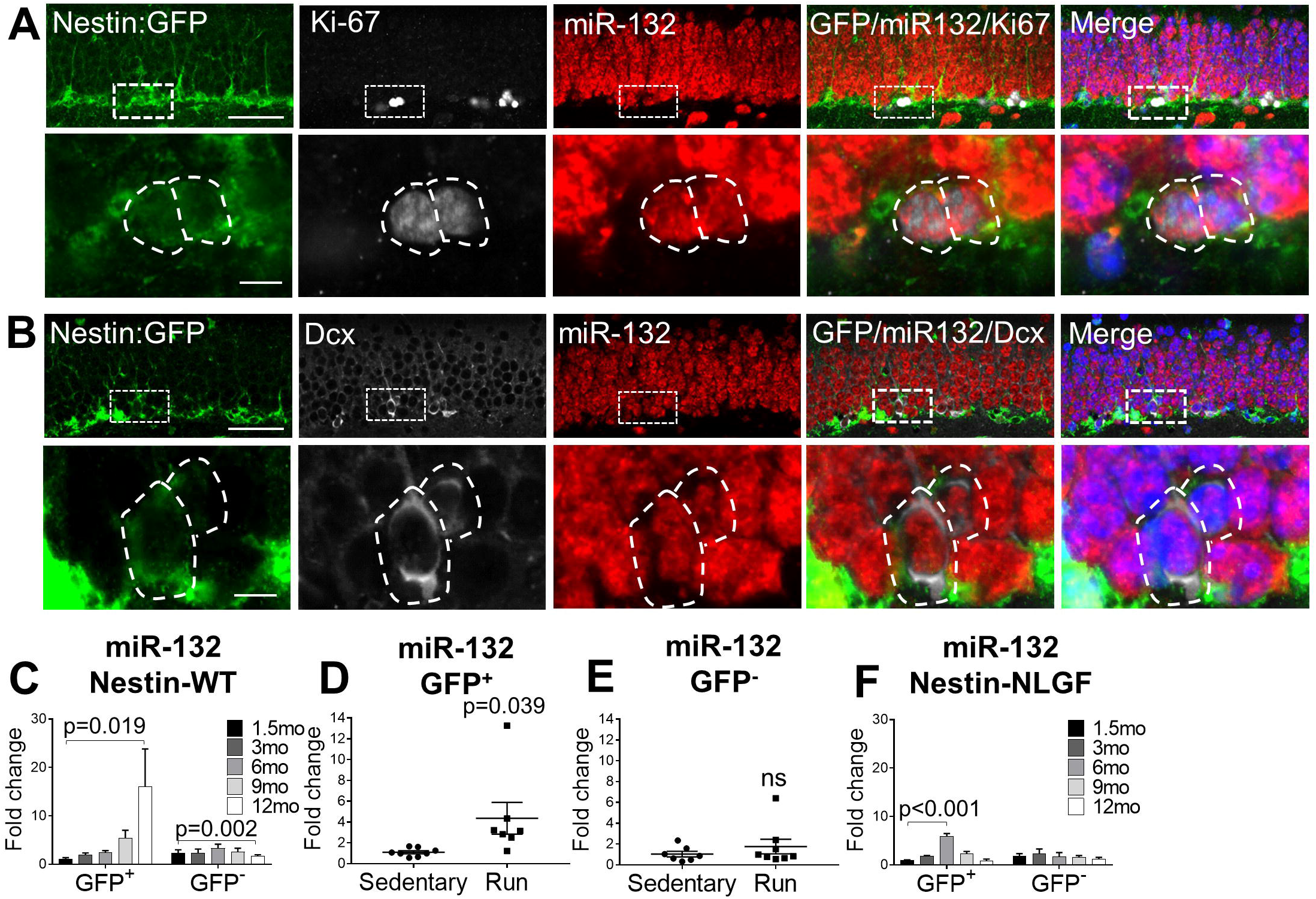
miR-132 in the adult hippocampal neurogenic niche and in cultured neural stem cells. A, B. miR-132 *in situ* hybridization coupled to GFP and Ki-67 (A) or GFP and doublecortin (DCX) (B) immunolabeling in the dentate gyrus of 3-month-old Nestin:GFP mice. Lower panels show magnified views of the cells in the regions indicated by dashed rectangles in upper panels. Dashed outlines in lower panels indicate cellular margins. Scale bars, 50 μm (upper panels); 10 μm (lower panels). C. Semi-quantitative real-time PCR of miR-132 levels in GFP^+^ or GFP^−^ populations isolated by FACS from the dentate gyrus of Nestin:GFP mice at 1.5, 3, 6, 9 and 12 months of age. N=6-8 mice per time point. D, E. Semi-quantitative real-time PCR of miR-132 levels in GFP^+^ (D) or GFP^−^ (E) populations sorted from the dentate gyrus of sedentary control or running Nestin:GFP mice at 3 months of age. N=6-8 mice per group. F. Semi-quantitative real-time PCR of miR-132 levels in GFP^+^ or GFP^−^ populations sorted from the dentate gyrus of Nestin-NLGF mice at 1.5, 3, 6, 9 and 12 months of age. N=3-5 mice per time point. Values are presented as mean ± SEM. In (C, F), two-way ANOVA with Tukey’s *post hoc* test for multiple comparisons were applied, while in (D, E), Student’s t-test was used. **See also Figure S2 and Table S4**.

### miR-132 is required for proliferation of neural precursors and neuronal differentiation in adult dentate gyrus

We then asked whether miR-132 is required for adult neurogenesis in wild-type dentate gyrus. As expected, voluntary running induced an increase in the numbers of BrdU^+^- and Ki-67^+^-proliferating neuronal precursors, Nestin:GFP^+^-neural stem cells and progenitors, and DCX^+^-immature neurons at the subgranular zone of the dentate gyrus in scramble control-injected wild-type mice compared to sedentary animals (Figures 3A–F). However, the effect of physical exercise on AHN was abrogated by miR-132 knockdown (Figure S3A) in running mice intracerebroventricularly (ICV)-injected with an antisense oligonucleotide against miR-132 (AntagomiR-132, Ant-132) (Figures 3A–F), demonstrating that miR-132 is required for the induction of AHN by running. miR-132 knockdown was confirmed in both the Nestin:GFP^+^- and the Nestin:GFP^−^ -fractions of the neurogenic niche (Figure S3B) and was specific to miR-132 (Figure S3C). Immunostaining against the proliferation marker PCNA confirmed that miR-132 knockdown abolishes the positive effect of running on neuronal precursor proliferation (Figures S3D, E). Since miR-132 is also involved in the regulation of neuronal apoptosis (Wong et al., 2013), we asked whether the net effect of miR-132 knockdown could be mediated by interfering with the physiological process of neuronal progenitor elimination by apoptotic death. Quantification of the cells labeled by an antibody against cleaved caspase-3, a marker of apoptosis, did not reveal any significant changes between groups (Figures S3F, G), indicating that miR-132 regulation of AHN is not effectuated at the level of apoptotic control. Interestingly, miR-132 deficiency also impaired the exercise-induced elevation of Brain-Derived Neurotrophic Factor (*Bdnf*) (Figure S3H), which has recently emerged as an essential neurotrophin for the fitness of the adult hippocampal neurogenic niche and the cognitive recovery in an AD mouse model (Choi et al., 2018).

**Figure 3.**
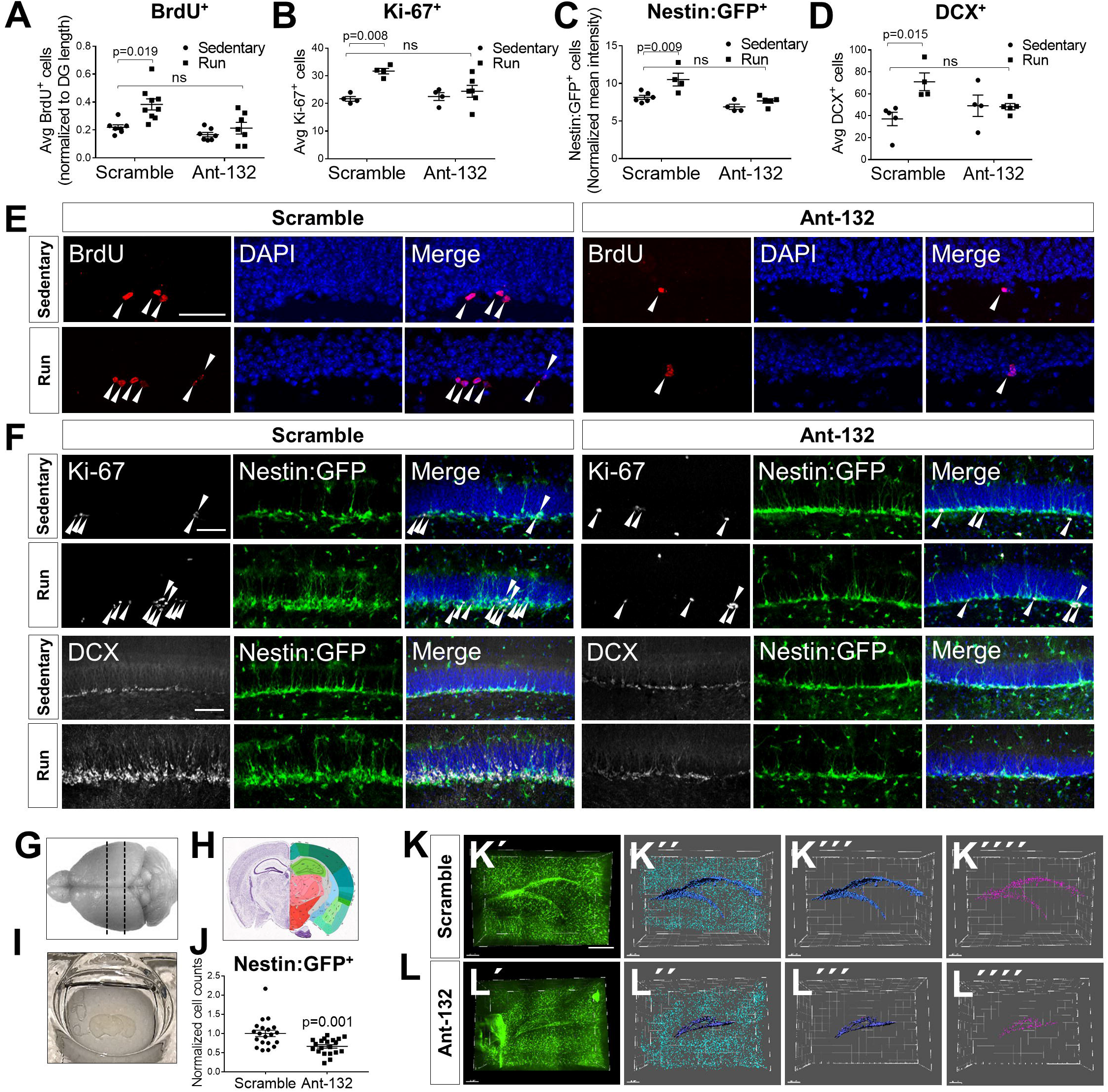
miR-132 is required for adult neurogenesis. A. Quantification of BrdU^+^-cells in the dentate gyrus of control- or AntagomiR-132 (Ant-132) -injected, sedentary or running C57Bl/6 mice at 9 months of age. N=6-9 mice per group. B-D. Quantification of Ki-67^+^-(B), Nestin:GFP^+^- (C) and DCX^+^- cells (D) in the dentate gyrus of control- or Ant-132-injected, sedentary or running Nestin:GFP mice at 3 months of age. N=4-6 mice per group. E. BrdU^+^-proliferating neuronal progenitors in the dentate gyrus of scramble- or Ant-132-injected, sedentary or running C57Bl/6 mice at 9 months of age. Arrowheads indicate BrdU^+^-cells. Scale bars, 50 μm. F. Ki-67^+^-proliferating progenitors and DCX^+^- immature neurons in control- or Ant-132-injected, sedentary or running Nestin:GFP mice at 3 months of age. Arrowheads indicate Ki-67^+^-amplifying progenitors. Scale bars, 50 μm. G. Schematic of mouse brain indicating the area of sectioning (dashed lines) used for tissue clearing. H. Coronal mouse brain section at P56, position 293, showing hippocampal structure. Image adapted from Allen Brain Atlas (http://mouse.brain-map.org/experiment/thumbnails/100048576?image_type=atlas). I. Cleared coronal brain section. J. Quantification of GFP^+^-cells in the subgranular layer of the dentate gyrus in running scramble- or Ant-132-injected 3-month-old Nestin:GFP mice. N=10 mice per group; 2 hemispheric dentate gyri per mouse. K, L. 3D reconstruction and image processing for whole-mount GFP^+^-cell counting in scramble- (K) or Ant-132- (L) injected mice upon running. Green, GFP^+^-cells; dark blue, reconstructed dentate gyrus surface; purple, GFP^+^-cells lining the dentate gyrus used for quantification; light blue, GFP^+^-cells away from dentate gyrus. Scale bars, 300 μm. Values are presented as mean ± SEM. Two-way ANOVA with Tukey’s *post hoc* test for multiple comparisons were employed in (A-D) and Student’s t-test in (J). **See also Figure S3**.

To obtain a measurement of the global effect of miR-132 deficiency on the neural stem cells and progenitors *in vivo*, we employed tissue clearing to assess the numbers of the Nestin:GFP^+^-cells at the subgranular zone of the whole dentate gyrus (Figures 3G–I). We compared the numbers of Nestin:GFP^+^-neural stem cells and progenitors at the subgranular zone of running mice (Figures 3J–L) injected either with scramble control- (Video S1) or with Ant-132 (Video S2). miR-132 knockdown resulted in a small but significant decrease in the total number of neural stem cells and progenitors (Figure 3J), confirming our previous observations. Together, these data unequivocally demonstrate that miR-132 is indispensable for the induction of neurogenesis the dentate gyrus *in vivo*.

### miR-132 regulates neuronal differentiation and is downregulated by AD-related pathology in human neural stem cells

We next assessed whether miR-132 regulation is also involved in the differentiation of human neuronal precursor to neurons. miR-132 levels significantly increased by 20-fold upon neural induction of primary human embryonic stem cells to neuronal precursors and were further boosted by more than 300-fold during neuronal maturation (Figures 4A, B). miR-212 levels, in contrast, were only subtly altered over the course of neuronal differentiation and maturation (Figure S4A). The progression of neuronal differentiation was monitored by measuring beta tubulin 3 (TUBB3) mRNA levels (Figure 4C).

**Figure 4.**
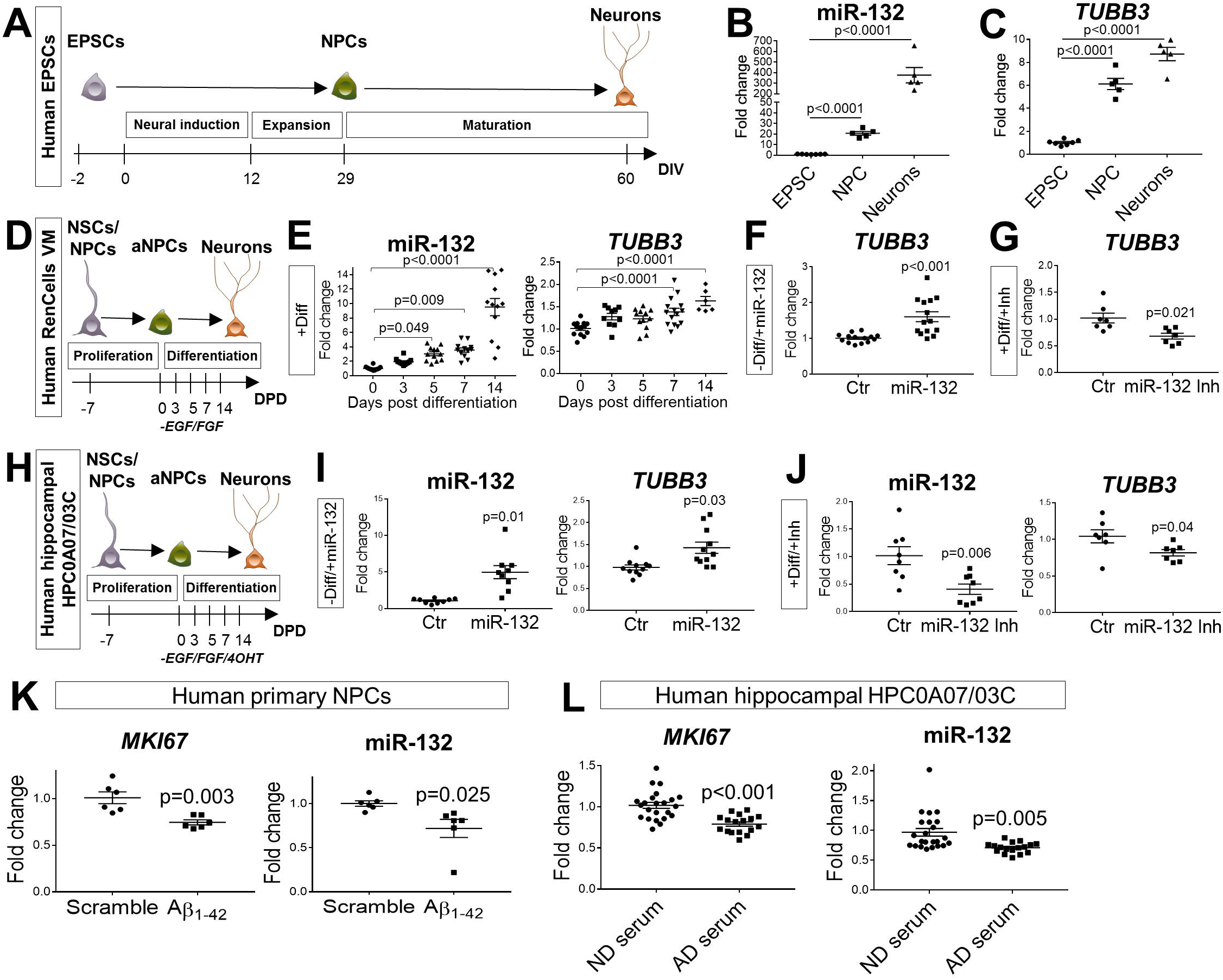
miR-132 regulatory effects in human neural stem cells. A. Schematic representation of the protocol used for neural induction of human embryonic pluripotent stem cells (EPSCs). DIV, days *in vitro*. B, C. Semi-quantitative real-time PCR of miR-132 (B) and tubulin (*TUBB3*) (C) levels in EPSCs (Day −2), NPCs (Day 29) and mature neurons (Day 60). N=3 independent experiments. D. Schematic diagram of the protocol used for neural differentiation of cultured immortalized human mesencephalic neuronal progenitors. NSCs/NPCs, neural stem cells/neuronal progenitor cells; aNPCs, amplifying neuronal progenitor cells; DPD, days post differentiation; E. Semi-quantitative real-time PCR of miR-132 and *TUBB3* upon induction of differentiation (Diff) in human neuronal progenitors. N=3 independent experiments. F, G. Semi-quantitative real-time PCR of *TUBB3* following miR-132 overexpression (miR-132) without induction of differentiation (F) or miR-132 knockdown (miR-132 Inh) upon induction of differentiation at 3 days post differentiation (G). N=3 independent experiments per condition. H. Schematic diagram of the protocol used for neural differentiation of cultured immortalized human hippocampal neuronal progenitors. NSCs/NPCs, neural stem cells/neuronal progenitor cells; aNPCs, amplifying neuronal progenitor cells; DPD, days post differentiation; I, J. Semi-quantitative real-time PCR of miR-132 and *TUBB3* following miR-132 overexpression (miR-132) without induction of differentiation (I) or miR-132 knockdown (miR-132 Inh) upon induction of differentiation at 3 days post differentiation (J). N=3 independent experiments per condition. K, L. Semi-quantitative real-time PCR of the proliferation marker *MKI67* (coding for Ki-67) and miR-132 in embryonic pluripotent stem cell-derived primary human neuronal precursor cells (NPC) (K) or an established human hippocampal neural precursor (hNPC) line (HPC0A07/03CI) (L) upon treatment with Aβ_1-42_ oligomers and a scramble control peptide (K) or incubation in AD and control serum (L). N=6 biological replicates in (K); 23 (ND) and 17 (AD) in (L). Values are presented as mean ± SEM. For data analysis, one-way ANOVA with Tukey’s *post hoc* test for multiple comparisons were applied in (B, C, E), while Student’s t-test was used in (F, G, I-L)). **See also Figure S4**.

A similar upregulation of miR-132 was detected in an immortalized human neural stem cell and progenitor line of mesencephalic origin (RenCells VM) upon induction of neuronal differentiation (Figures 4D, E), providing further support for the functional significance of miR-132 in the process of human neurogenesis. The differentiation efficiency was monitored by immunostaining for NESTIN and SOX2 (neural stem cells), Ki-67 (proliferating neuronal precursor cells) and *TUBB3* (mature neurons) (Figures S4B, C). The levels of *TUBB3* were additionally measured by semi-quantitative real-time PCR (Figure 4E). Notably, miR-132 transfection in human undifferentiated RenCell neural stem cells and progenitors (Figures S4D, E) resulted in a significant upregulation of *TUBB3* (Figures 4F, S4J), suggesting that miR-132 is a positive regulator of neurogenesis in human neuronal progenitors. A ten times lower miR-132 dose (Figure S4F) had an overall milder yet still significant effect on the levels of TUBB3 (Figure S4G). Conversely, miR-132 knockdown in differentiating RenCell neural stem cells and progenitors (Figures S4H, I) induced a decrease of TUBB3 levels (Figures 4G, S4K). Importantly, similar results were obtained upon miR-132 overexpression (Figure 4H, I) and knockdown (Figure 4J) in human neural precursor cells of hippocampal origin (HPC0A07/03C), corroborating the relevance of miR-132 regulation over human neurogenesis.

To explore whether Aβ pathology can prompt miR-132 downregulation in human neural stem and progenitor cells, we treated human embryonic stem cell-derived neuronal precursor cells with Aβ_1-42_ oligomers. Incubation of these cells with 5 μM Aβ_1-42_ for 72 h impacted their proliferation capacity (Figure 4K) and induced a specific miR-132 decrease compared to cells incubated with a scramble control (Figures 4K, S4L). The hippocampal neurogenic niche is highly vascularized and hence permissive to molecular cues transmitted via communication with the systemic environment (Villeda et al., 2011). Previously, it was shown that incubation of human neuronal precursor cells of hippocampal origin with serum samples derived from AD patients compared to non-demented control individuals reduced cell proliferation (Maruszak et al, 2017, BioRxiv 10.1101/175604). We extend this observation here by demonstrating that neuronal precursor cells treated with sera derived from AD patients showed significantly reduced miR-132, but not miR-212, levels (Figures 4L, S4M), while incubation with the AD sera also impacted cell proliferation (Figure 4L). These data confirm that miR-132 expression is suppressed in human stem cells by Aβ oligomers or other AD-related serum-derived factors.

### miR-132 alleviates proliferation and differentiation deficits of adult neural precursors in mouse AD brain

We next investigated whether boosting miR-132 levels could ameliorate the AHN deficits that we observed in the two AD mouse models. Control-injected 9-month old *APP/PS1* mice were not able to induce AHN upon running, as assessed by the numbers of BrdU^+^-, Ki-67^+^-proliferating neuronal precursor cells, and DCX^+^-immature neurons at the subgranular zone of the dentate gyrus (Figures 5A–D). However, upon miR-132 overexpression via ICV injection of a synthetic miR-132 mimic oligonucleotide (Figure S5A), we observed significant increases in these cell populations under sedentary conditions. Similar results were obtained in the *App^NL-G-F^* male (Figures 5E–H and S5B) and *App^NL-G-F^* female mice (Figures S5C, D), suggesting that this was not a strain-or gender-dependent effect. To assess whether the apparent increase in proliferating progenitors and immature neurons is the result of an intrinsic transcriptional response of the resident niche cells to miR-132 and to account for the variation in cell numbers between miR-132-overexpressing and control animals, we measured the levels of markers of stemness (*Pax6*), cell proliferation (*Mki67*) and early neuronal fate (Dcx) in equal numbers of sorted Nestin:GFP^+^-cells isolated from the dentate gyrus of miR-132- or control-injected mice (Figures 5I, S7A). miR-132 overexpression significantly repressed early stem cell identity in this pool of cells and induced transcriptomic changes indicative of increased proliferation and differentiation rates.

**Figure 5.**
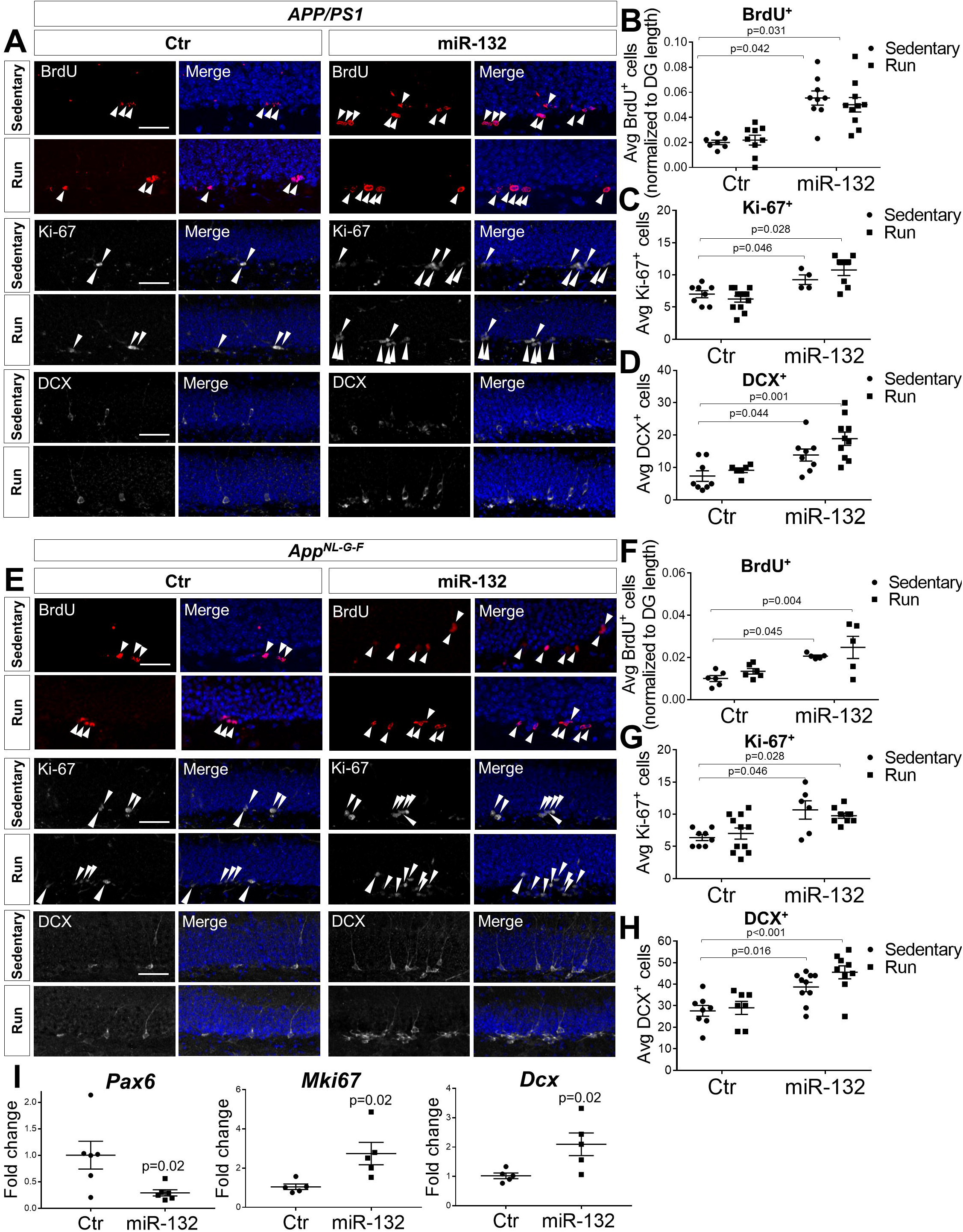
miR-132 overexpression ameliorates adult neurogenesis deficits in AD mouse hippocampus. A. Immunolabeling of BrdU^+^-, Ki-67^+^- and DCX^+^-cells in the dentate gyrus of control-(Ctr) or miR-132 (miR-132)-injected, sedentary or running *APP/PS1* mice at 9 months of age. Scale bar, 50 μm. B-D. Quantification of BrdU^+^- (B), Ki-67^+^- (C) and DCX^+^-cells (D) in the dentate gyrus of control- (Ctr) or miR-132 (miR-132)-injected, sedentary or running *APP/PS1* mice at 9 months of age. N=6-10 mice per group. E. Immunolabeling of BrdU^+^-, Ki-67^+^- and DCX^+^-cells in the dentate gyrus of control- (Ctr) or miR-132 (miR-132)-injected, sedentary or running *App^NL-G-F^* mice at 9 months of age. Scale bar, 50 μm. F-H. Quantification of BrdU^+^- (F), Ki-67^+^- (G) and DCX^+^-cells (H) in the dentate gyrus of control- (Ctr) or miR-132 (miR-132)-injected, sedentary or running *App^NL-G-F^* mice at 9 months of age. N=6-10 mice per group. I. Semi-quantitative real-time PCR of *Pax6*, *Mki67* and *Dcx* levels in GFP^+^-cells sorted from the dentate gyrus of Nestin:GFP mice at 3 months of age upon control- or miR-132-injection. N=5-6 mice per group. Values are presented as mean ± SEM. For data analysis, two-way ANOVA with Tukey’s *post hoc* test for multiple comparisons were applied in (B-D, F-H), while Student’s t-test was used in (I). **See also Figure S5**.

### Cell-autonomous regulatory effects of miR-132 in the adult neural stem cells

Considering that ICV infusion of miR-132 mimic or antisense oligonucleotides may exert broad effects in different cell types at the adult hippocampal niche, we used a lentivirus-mediated labeling approach to overexpress or knockdown miR-132 specifically in adult neural stem cells (NSC) and their progeny. In particular, lentiviral vectors (LV PGK:mCHERRY) expressing the mCHERRY reporter and either a miR-132 shRNA (for miR-132 knockdown) or a miR-132 hairpin (for miR-132 overexpression) all under the control of the PGK promoter (Suh et al., 2018), were used for stereotactical injections into the dentate gyrus (Figure 6A, B). Morphological development of the labelled cells followed a stereotypical temporal progression (Figure S6A). The efficiency of miR-132 knockdown or overexpression was assessed in FACsorted mCHERRY-positive cells derived from the dentate gyrus of mice injected with the shRNA, hairpin or corresponding control vectors (Figure 6C).

**Figure 6.**
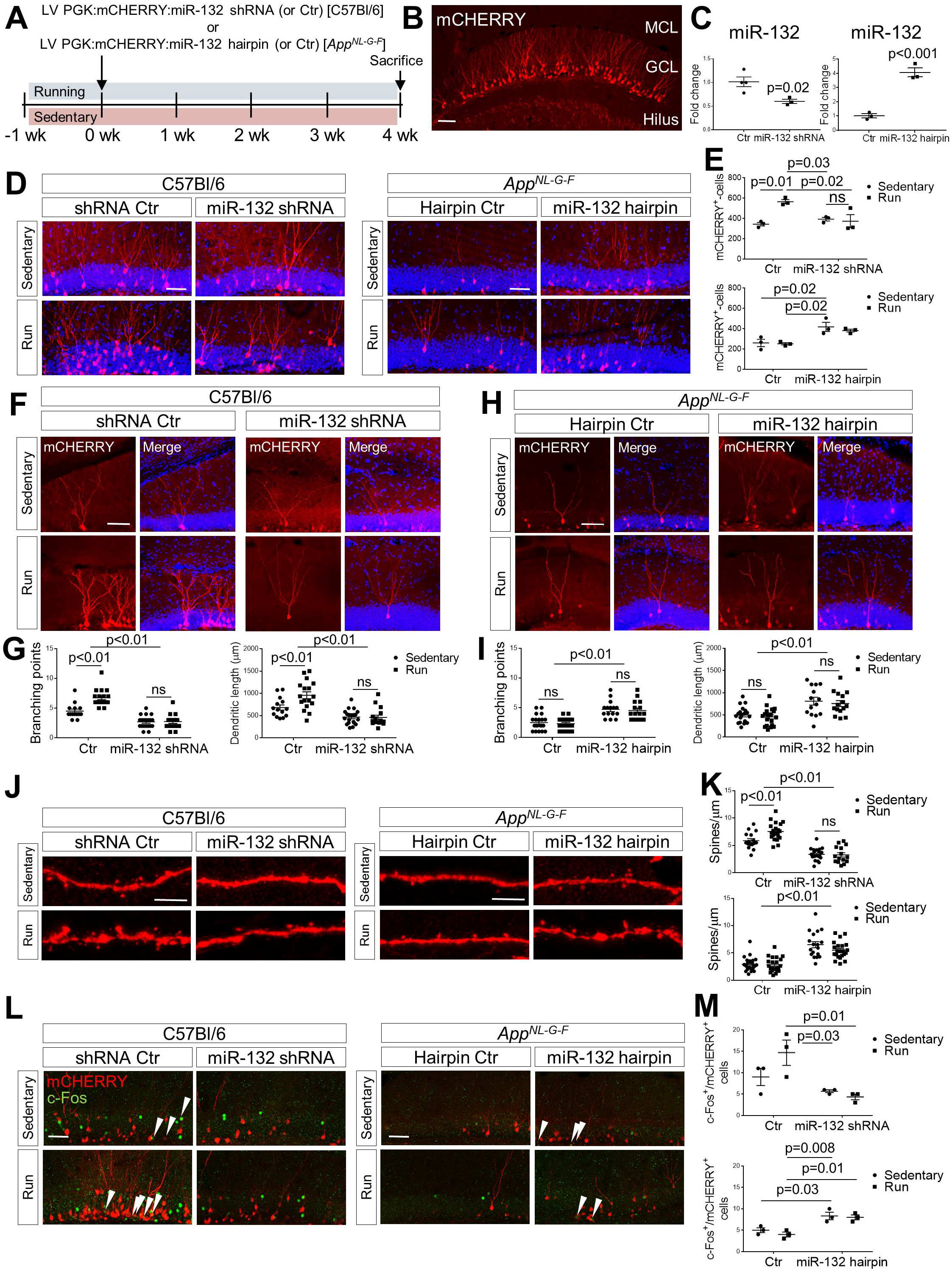
Cell-autonomous effects of miR-132 in adult-born granule neurons in wild-type and AD brain. A. Schematic diagram of the experimental design. Lentiviral vectors were used for stereotactical injections into the dentate gyrus of 2-month old C57Bl/6 or 9-month old *App^NL-G-F^* mice. Animals were housed under sedentary or running conditions and were euthanized 4 weeks post injection. B. Representative image of the mCHERRY-labeled 4-week old neurons in the dentate gyrus upon injection with the control lentiviral vector. Scale bar, 50 μm. C. Semi-quantitative real-time PCR of miR-132 in FACsorted mCHERRY^+^-cells upon miR-132 knockdown (miR-132 shRNA) or overexpression (miR-132 hairpin) compared to the respective control vector injections. N=3-4 mice per group. D. Immunolabeling of mCHERRY^+^- 4-week old neurons upon miR-132 knockdown in C57Bl/6 mice or miR-132 overexpression in *App^NL-G-F^* animals, under sedentary or running conditions. E. Quantification of the number of mCHERRY^+^- 4-week old neurons indicated in D. N=3 mice per group. F, G. Imaging and quantification of dendritic arborization in 4-week old neurons upon miR-132 knockdown in C57Bl/6 mice, under sedentary or running conditions. N=4-6 cells per mouse; 3 mice per group. H, I. Imaging and quantification of dendritic arborization in 4-week old neurons upon miR-132 overexpression in *App^NL-G-F^* animals, under sedentary or running conditions. N=4-6 cells per mouse; 3 mice per group. J, K. Imaging and quantification of spine density in 4-week old neurons upon miR-132 knockdown in C57Bl/6 mice or miR-132 overexpression in *App^NL-G-F^* animals, under sedentary or running conditions. N=1 dendrite per cell; 4-6 cells per mouse; 3 mice per group L, M. Imaging and quantification of the number of c-Fos^+^/mCHERRY^+^-cells upon miR-132 knockdown in C57Bl/6 mice or miR-132 overexpression in *App^NL-G-F^* animals, under sedentary or running conditions. Arrowheads indicate double-positive cells. Scale bars, 50 μm (B, D, F, H, L); 10 μm (J). Values are presented as mean ± SEM. For data analysis, two-way ANOVA with Tukey’s *post hoc* test for multiple comparisons were applied in (E, G, I, K, M), while Student’s t-test was used in (C,E). **See also Figure S6**.

Morphometric dendritic features in adult-born granule neurons are correlated to functional integration and connectivity (van Praag et al., 2002; Trinchero et al., 2017; Zhao, 2006). We analyzed dendritic development in 4-week old neurons, a time point at which these cells are functionally integrated (van Praag et al., 2002; Zhao, 2006). We confirmed that in wild-type animals, running increased the number and dendritic complexity of wild-type 4-week old granule cells, as indicated by the counts of mCHERRY-positive cells and the dendritic length and branches 4 weeks post injection (Figure 6D–G) (van Praag et al., 2005; Trinchero et al., 2017). Interestingly, selective miR-132 knockdown in the newly born neurons suppressed the running-induced increase in their population (Figure 5D, E). miR-132 knockdown also decreased dendritic arbor complexity in running but also in sedentary animals, suggesting a stimulation-independent, cell-autonomous effect of miR-132 on baseline dendritic development and maturation (Figure 6F, G). Consistent with the AHN deficits, the number and dendritic arborization of 4-week old granule neurons were lower in sedentary *App^NL-G-F^* mice compared to wild-type animals, and no running-induced effects were observed (Figure 6D, E, H, I). However, adult NSC-specific miR-132 overexpression reversed these deficits in the neuronal progeny under both sedentary and running conditions (Figure 6D, E, H, I).

Synaptic integration of the developing adult-born granule neurons follows a stereotypic sequence, starting with the formation of GABAergic dendritic inputs and followed by glutamatergic synaptogenesis and formation of dendritic spines (Trinchero et al., 2017). At 4 weeks of age, newly born neurons already receive excitatory input from the performant path, which connects the dentate gyrus to the entorhinal cortex (van Praag et al., 2002). While running increased dendritic spine density in wild-type control mice, lentiviral miR-132 knockdown in wild-type adult NSCs resulted in reduced spine density in 4-week old granule neurons (Figure 6J, K). Reversely, selective miR-132 overexpression in *App^NL-G-F^* adult-born neurons, restored spine density back to wild-type, as opposed to running, which had no effect (Figure 6J, K).

To monitor whether, along with the observed structural remodeling, miR-132 also exerts cell-autonomous regulatory effects on the ability of the adult-born neurons to receive excitatory input, we assessed the expression of c-Fos, an immediate-early gene, which has been widely used as a marker for neuronal activation (Clark et al., 2011). miR-132 knockdown resulted in a significant decrease in c-Fos/mCHERRY double-positive 4-week old neurons in running wild-type mice, while miR-132 overexpression in *App^NL-G-F^* adult-born neurons restored the number of double-positive cells back to wild-type (Figure 6L, M). Correction for the total amount of mCHERRY-positive cells (Figure S6B, C) retained the significant miR-132 knockdown-mediated impact on the c-Fos/mCHERRY double-positive 4-week old wild-type neurons, suggesting a putative cell-intrinsic inhibitory effect on excitability. Upon NSC-specific miR-132 knockdown or overexpression, we also observed significant changes in the non-mCHERRY-labeled, c-Fos^+^-cells in the granule cell layer. This suggests additional non cell-autonomous effects on the activity of other neurons exerted via the miR-132 knockdown or overexpression in the neural stem cells (Figure S6B, C). Finally, only very few Dcx/mCHERRY double-positive 4-week old neurons were detected, while the vast majority of the mCHERRY^+^-cells in all conditions were also NeuN-positive. No miR-132-dependent changes were detected in the number of Dcx/mCHERRY or NeuN/mCHERRY double-positive cells in neither wild-type nor *App^NL-G-F^* mice (Figure S6D-F).

Together, these data suggest that miR-132 regulates late-stage neurogenic events such as maturation and activation in a cell-autonomous manner, and that increasing miR-132 in adult neural stem cells can ameliorate deficits in the dentate gyrus of an AD mouse model.

### miR-132 promotes AHN by modulating a complex molecular network in adult neural stem cells

In order to identify the mechanisms underpinning the positive regulatory effect of miR-132 in AHN, we employed a single-cell RNA-sequencing approach to assess miR-132-specific transcriptomic responses in the Nestin:GFP-labeled resident niche cells (Figure 7A). Nestin:GFP^+^-niche cells were isolated from dentate gyri of mice that were ICV-injected with either miR-132 mimic or control oligonucleotides and single-cell libraries were prepared as discussed in the STAR Methods section. The extent and specificity of miR-132 overexpression was assessed in Nestin:GFP^+^ and Nestin:GFP^−^ bulk-sorted cells (Figure S7A). While miR-132 levels were, as expected, elevated in both populations compared to control-injected animals, the increase was only significant in the Nestin:GFP^+^ fraction (Figure S7A), which may reflect the relative high baseline expression of miR-132 in the neuronal fraction of the dentate gyrus. A total of 709 Nestin:GFP^+^ single niche cells were sequenced using the Smart-seq2 protocol. Unsupervised hierarchical clustering of the dataset using the Seurat algorithm revealed 7 populations, which were projected on a t-distributed stochastic neighbor embedding (t-SNE) map for visualization (Figure 7B). Based on the expression levels of well-characterized marker genes (Artegiani et al., 2017; Hochgerner et al., 2018; Zeisel et al., 2018) (Table S5; LiteratureMarkers), we defined cluster identity (Figures 7B, S7B and Table S5; ModuleScore), mapping the subpopulations as radial glia-like (RGL) neural stem cells, neuronal intermediate precursor cells (NPC), astrocytes, oligodendrocyte precursor cells (OPC), myelin-forming oligodendrocytes (MFOL), endothelial cells and pericytes. Cell type-enriched genes were identified by comparing each cell type cluster to all others (Figure S7C and Table S5; ClusterMarkers). The distribution of cells across clusters was not altered in a statistically significant way upon miR-132 overexpression in these experiments (data not shown).

**Figure 7.**
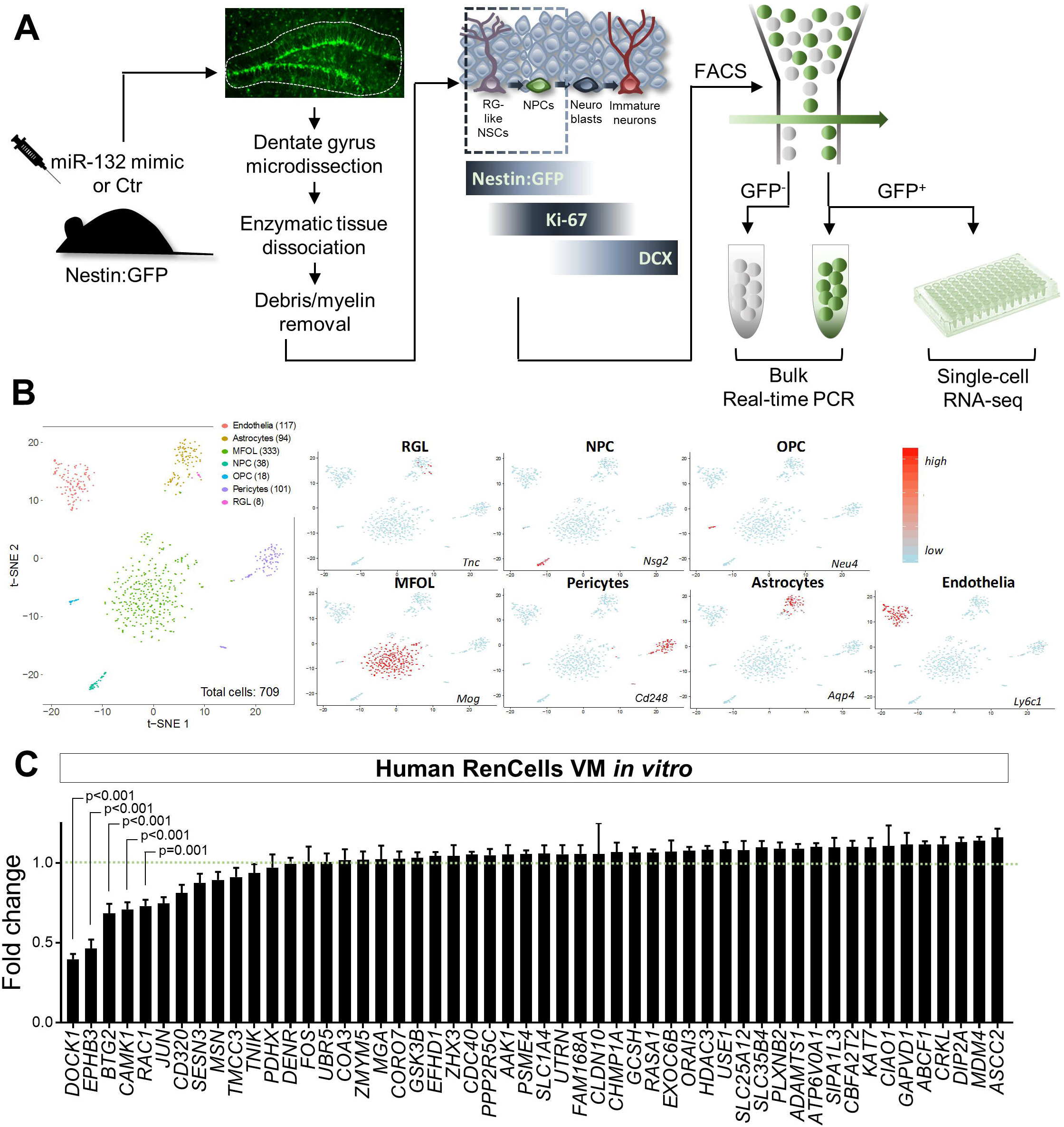
Single-cell approach to identify miR-132 targets in AHN. A. Schematic diagram of the experimental workflow used to isolate and analyze adult Nestin:GFP^+^ niche cells from the mouse dentate gyrus upon miR-132 overexpression. B. Unsupervised hierarchical clustering of dataset, cell type mapping and cell type-specific marker expression. Cell numbers per cluster are indicated in parentheses. C. Semi-quantitative real-time PCR assessment of predicted miR-132 target transcripts in the human RenCell line upon transfection with miR-132 mimic or a control oligonucleotide. N=3 independent experiments. RGL, radial glia-like cells; NPC, neuronal intermediate progenitor cells; OPC, oligodendrocyte precursors; MFOL, myelin-forming oligodendrocytes; In C, two-way ANOVA with Tukey’s *post hoc* test for multiple comparisons were used. Dashed line indicates mean fold change of control samples set at 1. Values are presented as mean ± SEM. **See also Figure S7 and Tables S5, S6**.

Differential gene expression analysis between miR-132-overexpressing and control cells within each cluster did not reveal any significantly deregulated transcripts after correction for multiple testing comparisons (Table S6), an observation that could be indicative of the frequently documented limited effect sizes of microRNA regulation (Moore et al., 2015). Since microRNAs may elicit changes on cellular dynamics via widespread, yet subtle effects on the transcriptome (Liufu et al., 2017), we next assessed the possible impact of miR-132 overexpression on molecular pathway regulation through gene set analysis. Gene ontology (GO) enrichment analysis using Ingenuity Pathway Analysis (IPA, Ingenuity Systems Inc., Redwood City, CA, USA) identified significantly enriched pathways upon miR-132 overexpression in each of the seven cell subpopulations (Figure S7D), suggesting that miR-132 regulation, although subtle at the single-gene level, may concomitantly impact diverse cellular processes. To gain empirical support for this notion, we used the human RenCell line to validate putative miR-132 targets. To prioritize possible direct targets, we selected 150 of the most downregulated transcripts in single neural stem cells (radial glia-like cells) upon miR-132 overexpression. Using three prediction algorithms for miRNA target identification (Targetscan, DIANA, PicTar), we extracted a list of 52 predicted miR-132 targets (predicted at least by one of the three algorithms) for semi-quantitative real-time PCR validation in RenCells transfected with either a miR-132 mimic or a control oligonucleotide. Interestingly, there was a significant change in the levels of five mRNAs i.e. *DOCK1* (Dedicator Of Cytokinesis 1), *EPHB3* (*EPHrin* receptor B3), *BTG2* (B-cell Translocation Gene anti-proliferation factor 2), *CAMK1* (Calcium/Calmodulin-Dependent Protein Kinase 1) and *RAC1* (RAs-related C3 botulinum toxin substrate 1), which were all found to be downregulated (Figure 7C), suggesting that these transcripts may be part of the direct effectors of the miR-132-regulated molecular network that generates proneurogenic signals in the radial glia-like neural stem cells.

### Boosting miR-132 levels restores AHN-related memory deficits in old AD mice

AHN has been linked to the memory processing ability of the hippocampal circuitry and to several mnemonic processes, including contextual memory and, in particular, pattern separation (Clelland et al., 2009; Suárez-Pereira et al., 2015). To address the functional impact of AHN regulation by miR-132 on relevant memory functions, we assessed the effect of miR-132 overexpression in 9-month old *App^NL-G-F^* mice in a passive avoidance test and in a pattern separation, differential fear conditioning-based test (van Boxelaere et al., 2017). Age-matched wild-type mice were used as controls and animals were ICV-injected with either a negative control oligonucleotide or a synthetic miR-132 mimic before behavioral testing (Figure S5E). In passive avoidance, AD mice exhibited decreased latency time to enter the conditioning chamber compared to wild-type animals during the testing phase, suggesting impaired contextual memory (Figure 8A). No change was observed between controls and miR-132-overexpressing wild-type mice (Figure 8A). Interestingly, boosting miR-132 levels in the AD mice rescued this deficit and restored latency time to wild-type levels (Figure 8A). In contrast, after repeated contextual conditioning in a pattern separation test (Figures 8B, C), AD mice showed similar contextual fear memory as wild-type animals, and elevating miR-132 levels had no effect on contextual discrimination between two very distinct contexts (Figure 8D, contexts A and C). These results indicate that repeated conditioning establishes context-specific memory traces also in the AD animals, while boosting miR-132 levels has no effect on specific contextual memory. However, discriminating between highly similar contexts (contexts A and B) relies on pattern separation, which is dependent upon the firing properties of the adult-born neurons in the dentate gyrus. In this test, wild-type controls displayed discriminatory levels of freezing in contexts A and B, while AD animals were unable to distinguish between the two overlapping contexts (Figure 8E). Interestingly, miR-132 overexpression in *App^NL-G-F^* mice restored performance in the pattern separation test (Figure 8E), suggesting that the positive regulation of AHN by miR-132 can promote cognitive recovery in AD. However, overexpressing miR-132 in wild-type mouse brain resulted in poor performance compared to the control-injected group (Figure 8E), which could indicate a possible requirement for a dose window for the miR-132 positive effect.

**Figure 8.**
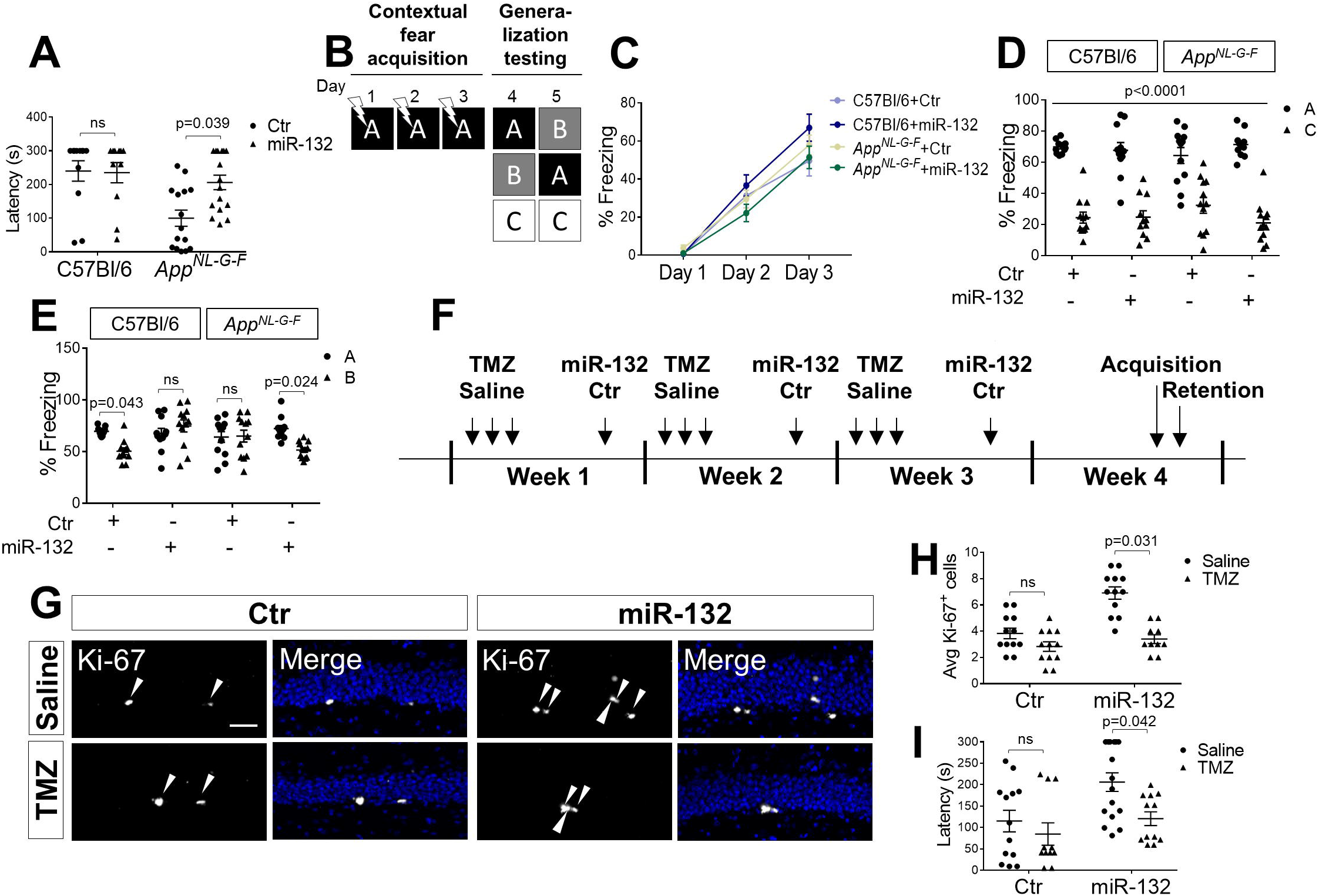
miR-132 overexpression restores memory deficits in AD mice. A. Latency time (indicated in seconds, s) in passive avoidance test upon control or miR-132 injections in wild-type and AD mice. B. Schematic diagram of pattern separation test. Mice were fear-conditioned for 3 days in shock-receiving context A, and subsequently tested for their ability to discriminate between similar contexts A and B, and between different contexts A and C. C. Percentage of time spent freezing by control (Ctr)- or miR-132 (miR-132)-injected 9-month-old wild-type (C57Bl/6) or *App^NL-G-F^* mice during days 1-3 of fear acquisition. D, E. Percentage of time spent freezing in contexts A and C (D) or A and B (E) during days 4-5 of generalization testing. N=12-15 mice per group. F. Schematic diagram of passive avoidance test in 9-month-old *App^NL-G-F^* mice employing saline or temozolomide (TMZ) intraperitoneal (i.p.) injections and control (Ctr) or miR-132 (miR-132) intracerebroventricular injections. G. Ki-67 immunolabeling in the dentate gyrus of saline or TMZ-, Ctr- or miR-132-injected *App^NL-G-F^* mice after passive avoidance test completion. H. Quantification of Ki-67^+^-cells upon saline or TMZ injections in Ctr- or miR-132-injected *App^NL-G-F^* mice. N=11-13 mice per group. Scale bar, 50 μm. I. Latency time in passive avoidance test in saline or TMZ-injected, Ctr- or miR-132-injected *App^NL-G-F^* mice. Values are presented as mean ± SEM. N=11-13 mice per group. For data analysis, two-way ANOVA with Tukey’s *post hoc* test for multiple comparisons were employed. **See also Figure S5**.

We then addressed the possible contribution of AHN to the beneficial effect of miR-132 in the passive avoidance test in the AD mice. We used the DNA-alkylating agent temozolomide (TMZ), which has been previously shown to repress AHN and context discrimination (Egeland et al., 2017; Garthe et al., 2009; McAvoy et al., 2016; Niibori et al., 2012), to block cell proliferation in control- and miR-132-injected *App^NL-G-F^* mice (Figure 8F). Ki-67^+^- and Dcx^+^- cell counts at the subgranular zone of the dentate gyrus were used as a readout of the efficiency of proliferation inhibition of adult NPCs (Figures 8G, H, S5F, G). Blocking NPC proliferation in miR-132-overexpressing AD mice abolished memory rescue in the passive avoidance task (Figure 8I), indicating that cell proliferation is crucial, thereby providing indirect evidence that AHN may be a prime component of miR-132 regulation on contextual memory performance in this task. These results suggest that replacing miR-132 levels in the AD brain can restore AHN-related memory deficits.

## Discussion

Downregulation of miR-132 expression is a consistent observation in brain samples of Alzheimer’s disease patients (Hebert et al., 2013; Lau et al., 2013; Patrick et al., 2017; Pichler et al., 2017; Salta and De Strooper, 2017; Salta et al., 2016; Smith et al., 2015; Wong et al., 2013; Zhu et al., 2016). We demonstrate here significant reduction of miR-132 levels in the adult hippocampal neurogenic niche in two different amyloidosis mouse models for AD and confirm downregulation of miR-132 in two different *in vitro* models for human neuronal stem cell differentiation upon treatment with synthetic Aβ oligomers or serum from AD patients. We link these defects to impaired AHN in the two AD mouse models by showing that intracerebroventricular infusion of synthetic miR-132 oligonucleotides or lentivirus-based overexpression of miR-132 directly in adult neuronal stem cells rescues the AHN deficits and additionally restores aspects of memory dysfunction. Our work suggests that loss of miR-132 is an important contributor to the overall decreased AHN in Alzheimer’s patients (Moreno-Jiménez et al., 2019; Tobin et al., 2019) and provides evidence that restoring miR-132 levels in the adult hippocampal neurogenic niche might be beneficial in AD.

Whether AHN is affected in AD brain and, even more fundamentally, whether AHN occurs in adult and aging human brain at all, has been a returning issue of controversy (Boldrini et al., 2018; Eriksson et al., 1998; Gatt et al., 2019; Kempermann et al., 2018; Knoth et al., 2010; Lucassen et al., 2019, 2020; Paredes et al., 2018; Snyder, 2019; Sorrells et al., 2018; Spalding et al., 2013; Tartt et al., 2018). The discussion has taught the paramount importance of accurate interpretation of histological findings, sample stratification (to minimize confounding factors such as agonal state, coexisting brain pathology, neuroinflammation, medication), postmortem delay, tissue preservation, marker labeling methodology, and inclusion of stereology during data analysis (Kempermann et al., 2018; Lucassen et al., 2019, 2020). Notably, two recent reports using high-quality human postmortem brain samples and golden standard methodological approaches have now firmly established that adult neurogenesis occurs in human brain. These studies demonstrated that AHN can still be detected in centenarians, and that AHN drops dramatically in AD patients (Flor-García et al., 2020; Moreno-Jiménez et al., 2019; Tobin et al., 2019), setting the stage for the functional studies in two mouse models of AD reported here.

We show that the running-induced miR-132 upregulation in the hippocampal neurogenic niche becomes compromised with pathology progression, independently of gender, and that these alterations are paralleled by decreased neurogenic potential. The hippocampal neurogenic niche involves a complex network of intercellular communication and molecular regulatory signals, which is receptive to extrinsic and intrinsic cues, such as experience, exercise, aging and increased neuroinflammation (Fuster-Matanzo et al., 2013; Mosher and Schaffer, 2018; Villeda et al., 2011). Our data demonstrate that treatment of human neuronal precursor cells with oligomeric Aβ reduces proliferation and miR-132 expression, consistently with several lines of evidence suggesting that Aβ *per se* impacts the neurogenic potential of neural stem cells and progenitors (Haughey et al., 2002; He et al., 2013; Lee et al., 2013). Serum obtained from sporadic AD cases also elicited a significant reduction of miR-132 in hippocampal human neural stem cells. This can be attributed to Aβ oligomers or other AD-related systemic signals.

In an attempt to identify relevant transcriptomic targets of miR-132 at the adult hippocampal neurogenic niche, we performed single-cell transcriptomic analysis of the Nestin:GFP-positive niche-residing cells, isolated from the dentate gyrus of mice overexpressing miR-132. Although these studies revealed trends for regulation of transcripts in several cell types, these changes did not reach statistical significance. The inherent challenges of characterizing direct microRNA targets is well acknowledged (Liufu et al., 2017) and likely reflect the subtle pleiotropic roles of miR-132 in the adult neurogenic niche. Given these limitations, we decided to validate possible regulatory interactions in human neural stem cells *in vitro*. This approach revealed a novel miR-132-specific molecular signature consisting of *DOCK1, EPHB3, BTG2, CAMK1* and *RAC1*, all of which have reported roles in neuronal differentiation and function (Fortin et al., 2010; Goold and Nicoll, 2010; Pillat et al., 2016; Theus et al., 2010; Vadodaria et al., 2013; Yang et al., 2012). Dock1, Ephb3, Camk1 and Rac1 have additionally been implicated in AD (Aguilar et al., 2017; Becker et al., 2014; Overk and Masliah, 2014; Riascos et al., 2014), while all five proteins have been associated with the immune response in a variety of cells (D’Ambrosi et al., 2014; Goold and Nicoll, 2010; Guo et al., 2013; Naskar et al., 2014; Pillat et al., 2016; Theus et al., 2010; Yuniati et al., 2018). Interestingly, miR-132 knockdown also repressed the running-induced increase of *Bdnf*, a key neurotrophic factor contributing to the fitness of the niche and the neurogenic potential. Collectively, these observations suggest that the proneurogenic effects of miR-132 may be effectuated at the level of NPC proliferation, neuronal differentiation, survival, functional integration, neurotrophic boost or a combination thereof, possibly depending on the presence or absence of distinct stimuli.

Adult-born neurons can modulate and sparsify activity in the dentate gyrus. They act as autonomous coding units with unique electrophysiological properties, thereby enabling the hippocampus to optimize memory resolution and robustness (Johnston et al., 2016). These newly born cells facilitate the spatiotemporal contextualization of information and they help avoid catastrophic interference in the hippocampal network, by promoting ‘behavioral pattern separation’ (Kempermann et al., 2018). They may also contribute directly or indirectly to forgetting and memory clearance, memory consolidation and spatiotemporal contextualization (Gonçalves et al., 2016). Poor performance in pattern separation tasks in AD patients (Ally et al., 2013; Wesnes et al., 2014; Yassa et al., 2010) and preserved AHN in non-demented individuals with AD or MCI neuropathology have been reported (Briley et al., 2016; Tobin et al., 2019). While a comprehensive analysis of the relationship between miR-132, deficient AHN and memory would require a separate study, we confirm here that features of hippocampus-dependent functions that are related to context fear conditioning and avoidance tasks, which are linked to deficient AHN in the context of an AD mouse model, can be rescued by restoring miR-132 levels in the brain of these mice.

Intriguingly, miR-132 overexpression in wild-type mice resulted in a worsening of performance in the AHN-specific pattern separation task, suggesting that miR-132 expression should be maintained within a certain range to ensure proper learning and memory function. Indeed, while miR-132 levels are approximately 1.5-fold induced in mouse hippocampus upon presentation with a spatial memory task (Hansen et al., 2013), higher, supra-physiological (>3-fold) levels inhibit hippocampus-dependent memory in wild type mice (Hansen et al., 2013). Our approach yielded a miR-132 overexpression of approximately 4-fold in the dentate gyrus of old wild-type and AD mice, which improved the memory deficits in the AD mice but interfered with learning and memory in the wild-type animals. Thus, further work is required to assess a therapeutically relevant window for miR-132 overexpression in AD brain.

In conclusion, our data support a model where miR-132 acts as a proneurogenic signal transducer contributing to specific aspects of memory formation. AD pathology leads to miR-132 deficiency and ultimately compromises AHN. Recently, induction of AHN in conjunction with improvement of the microenvironment of the adult hippocampal niche via BDNF overexpression was put forward as a putative target for therapeutic intervention in AD (Choi et al., 2018). Moreover, recent groundbreaking advances in the treatment of neurodegenerative disorders using small RNA-based therapeutics in spinal muscular atrophy (Finkel et al., 2016) may also pave the way for novel miR-132-based therapeutic approaches in AD. Dose determination will be crucial, while delineating the molecular and cellular mechanisms underlying the neurogenic effects of miR-132 may provide impetus for identifying novel strategies to therapeutically harness AHN in AD.

## Acknowledgements

We thank Véronique Hendrickx and Jonas Verwaeren for animal husbandry; Sebastian Munck, Nikky Corthout, Axelle Kerstens (LiMoNe, VIB), the VIB Nucleomics Core of Leuven, the VIB-KU Leuven FACS Core, An Snellinx, Isabel Salas, Dries T’Syen, Sara Calafate, Jeroen Vanderlinden, Natalia Gunko, Dorien Vandael and Jenny Peeters for providing technical assistance and resources; Annerieke Sierksma, Patrik Verstreken, Laure Bally-Cuif, Joris De Wit, Pierre Vanderhaeghen and Matthew Holt for critical discussions and feedback. The *APP/PS1* mice were a gift from Mathias Jucker, DZNE, Germany, the *App^NL-G-F^* mice were kindly provided by Takaomi Saido, RIKEN Brain Science Institute, Japan, and the Nestin:GFP mice from Grigori Enikolopov, Stony Brook University, NY, USA. The PGK lentiviral vector was a gift from Antonella Consiglio. Confocal microscope equipment was acquired through a Hercules Type 1 AKUL/09/037 to Wim Annaert and a CLME grant to the VIB BioImaging Core. ES is supported by the Fonds voor Wetenschappelijk Onderzoek (FWO), the Alzheimer’s Association (AA) and the Alzheimer’s Research Foundation (Stichting Alzheimer Onderzoek, SAO). Work in the De Strooper laboratory is supported by a European Research Council (ERC) grant ERC-2010-AG_268675, FWO, KU Leuven, VIB, SAO, the Elisabeth Foundation, VIND (IWT 135043), and a Methusalem grant from KU Leuven and the Flemish Government. BDS is the Bax-Vanluffelen Chair for Alzheimer’s Disease and is supported directly by the Opening the Future campaign of the Leuven Universiteit Fonds (LUF). DRT is funded by grants from FWO (G0F8516N Odysseus) and Vlaamse Impulsfinanciering voor Netwerken voor Dementie-onderzoek (IWT 135043). The work in the Thuret lab is supported by UK Medical Research Council grants MR/N030087/1 and MR/S00484X/1.

## Author contributions

Conceptualization, ES and BDS; Methodology, ES, KC, LW, SB, ZCV, RDH, DRT, ESi, ST, CSF, BDS; Investigation, ES, HW, EVE, SS, KC, KH, LW, SB, AR, ESi, CSF; Software, MF, NT, YF; Formal Analysis, ES, MF, NT, YF; Resources, ZCV, RDH, DRT, HZ, ST; Writing – Review & Editing, ES and BDS, with input from all authors; Supervision, ES, BDS; Funding Acquisition, ES and BDS.

## STAR Methods

### Animals

The wild-type mice we used were C57BL/6. The AD mouse models used in the study were the *APP/PS1* and *App^NL-G-F^* strains. The first mouse model co-expresses human-mutated APP^Swe^ (KM670/671NL APP) and human mutated presenilin 1 (L166P). These mice [Tg(Thy1-APPSw, Thy1-PSEN1*L166P) 21Jckr] typically show amyloid deposition in the hippocampus at 3–4 months and cognitive impairment at 7–8 months of age (Radde et al., 2006; Serneels et al., 2009). *App^NL-G-F^* mice are a more recently established App knock-in model that expresses the APP KM670/671NL (Swedish), APP I716F (Iberian), APP E693G (Arctic) mutations. Similarly to the *APP/PS1* mice, *App^NL-G-F^* animals develop plaques around the age of 3 months and behavioral deficits from 6 months onwards (Saito et al., 2014). For neural stem cell and neuronal progenitor cell visualization *in vivo* and cell isolation from the adult dentate gyrus, the previously characterized Nestin:GFP mice were used (Mignone et al., 2004). For Nestin:GFP^+^-niche cell sorting from the AD dentate gyrus, Nestin:GFP mice were crossed to *App^NL-G-F^* animals (Nestin-NLGF). All animal experiments were approved by the ethical committees of KU Leuven and UZ Leuven (LA1210596).

### Human samples

The clinical and histopathological information regarding the human hippocampal samples used in the study is summarized in Table S1. Information on collection and processing of the serum samples is provided in Table S4. All experimental procedures with human samples were approved by the ethical committees of KU Leuven, UZ Leuven (S59654) and the University of Gothenburg.

### Intracerebroventricular (ICV) injections

The intracerebroventricular injections were performed as previously described (Jimenez-Mateos et al., 2011) using the following stereotactic coordinates: AP-0.1 mm, ML-1.0 mm, and DV-3.0 mm (from the skull). For miR-132 knockdown, 2 month-old Nestin:GFP or 8 month-old C57Bl/6 mice were infused with 2 μl of miR-132 antagomiR (locked nucleic acid (LNA)-, 3⍰-cholesterol-modified oligonucleotide) (Exiqon, Qiagen, Denmark) at 0.33 nmole/μl in artificial cerebrospinal fluid (aCSF) (Harvard Apparatus, USA). Control mice received a scrambled LNA oligonucleotide in aCSF. Mice were exposed to voluntary running one week post operation. Analysis of antagomiR-132-injected animals was performed at 3 and 9 months of age, respectively (after 4 weeks of voluntary running). For miR-132 overexpression, 2 month-old Nestin:GFP or 8 month-old *App^NL-G-F^* and *APP/PS1* mice received either a miR-132 mimic or a negative control oligonucleotide (Dharmacon, Horizon Discovery, Belgium) in a 3 μl mix with lipofectamine 2000 (at a 1:1 volume ratio) (Thermo Fischer Scientific, Belgium). Injections of 150 pmol oligo each were performed once a week for 5 weeks in total. One week after the first injection, mice were exposed to voluntary running for a total of 4 weeks. Analysis of miR-132 mimic-injected animals was carried out at 9 months of age. Randomization of injectates was employed for all injection sessions, and animals were randomly allocated to each treatment.

### Lentiviral vector injections into the dentate gyrus

Male and female C57Bl/6 (2-month old) or *App^NL-G-F^* (9-month old) mice were housed in standard (sedentary conditions) or with a running wheel 1 week prior to surgery. For miR-132 overexpression or knockdown specifically in adult neural stem cells, a lentiviral Puro:mPGK:mCHERRY backbone vector was used (Suh et al., 2018) harboring a miR-132 hairpin or miR-132 shRNA sequence, respectively (VectorBuilder, Germany). Hairpin or shRNA constructs corresponding to cel-miR-67 were used as negative controls. For stereotactic injections, virus was infused (2⍰μl/hemisphere at 0.5⍰μl⍰min^−1^) bilaterally into the right and left dentate gyrus (anteroposterior: −2 mm from Bregma; mediolateral: ±1.5 mm; dorsoventral: −2 mm) (van Praag et al., 2002). Mice were kept under sedentary or running conditions and were sacrificed 4 weeks post injection.

### Induction of AHN via voluntary running

Mice were housed individually in standard rat cages (45 cm × 20 cm × 20 cm) and divided into sedentary or running groups. Runners had unlimited access to a running wheel in their cage for 30 days. Running distance (km) was monitored electronically (MAFAC, France). No differences in running velocity or distance were observed between wild-type and AD animals.

### 5-Bromo-2’-deoxyuridine (BrdU) injections

BrdU (Sigma-Aldrich, Belgium) was dissolved in 0.9% NaCl at a concentration of 10 mg/ml and was filtered (0.2 μm) under sterile conditions. Mice were pulsed with a single i.p. injection of BrdU (50 mg/kg of body weight) daily for 3 days (at the end of the running period). 24 h after the last BrdU injection they were perfused with 4% paraformaldehyde (PFA) and processed for BrdU immunostaining to identify proliferating neuronal progenitors.

### Immunofluorescence in mouse brain sections and cultured cells

For coronal brain sections (16 μm-thick cryosections for VGlut1/PSD95 stainings; 20 μm-thick cryosections for BrdU staining; 40 μm-thick vibratome-prepared sections for Nestin:GFP, Ki-67, DCX staining), tissue was initially permeabilized in 1% (v/v) TritonX-100 (cryosections were first post-fixed in 4% PFA for BrdU and in acetone/methanol for VGlut1/PSD95) and then blocked in 1% (v/v) TritonX-100, 10% (v/v) normal goat serum in PBS for 2 h at room temperature. Primary antibody incubation was carried out in 0.3% (v/v) TritonX-100, 3% (v/v) normal goat serum overnight at 4°C followed by incubation in secondary antibody for 2 h at room temperature. Finally, sections were incubated in DAPI (Sigma-Aldrich, Belgium) and mounted in Mowiol. For immunofluorescent labeling in cells cultured on coverslips, samples were initially permeabilized like before and then incubated in blocking solution (1.5% (v/v) normal donkey serum, 0.2% (v/v) TritonX-100 in PBS) for 1 h at room temperature. Primary and secondary antibody incubation was carried out in blocking solution at 4°C overnight and for 2 h at room temperature, respectively. Finally, cells were stained with DAPI and mounted as described earlier. All the antibodies used for immunolabeling of brain tissue and cultured cells are listed in Table S2.

### Immunostaining of human hippocampal sections

An automated staining protocol (Bond Polymer Refine Detection Protocol, Leica Microsystems, Belgium) was followed using a Leica Autostainer (Leica Microsystems, Belgium). Briefly, following dewaxing and rehydration, 5 μm-thick formalin-fixed, paraffin-embedded hippocampal sections were first blocked in 3-4% (v/v) H_2_O_2_ and then incubated sequentially with the primary antibody for 30 min, with anti-rabbit poly-HRP IgGs, DAB (3,3’-Diaminobenzidine tetrahydrochloride hydrate) and hematoxylin. In the case of double stainings, the second primary antibody was added to the sections after hematoxylin staining and was followed by incubation in an AP-conjugated antibody and in Bond Polymer Refine Red Detection reagent (Leica Microsystems, Belgium). Finally, the sections were dehydrated and mounted on a Leica automated Coverslipper (Leica Microsystems, Belgium). The antibodies used for immunostaining of human brain samples are listed in Table S2.

### Tissue clearing

For brain clearing, the CUBIC protocol (Susaki et al., 2015) was adapted. Briefly, two 1 mm-thick coronal brain sections (Interaural: 1.68/0.88 mm, Bregma: −2.12/-2.92 mm) from 4% PFA-perfused brains were initially immersed in 12.5% (w/w) urea, 12.5% (w/w) quadrol and 7.5% (w/w) TritonX-100 for 6 h at 37°C for lipid removal. Subsequently, samples were incubated in 25% (w/w) urea, 25% (w/w) quadrol, and 15% (w/w) TritonX-100 for 24 h at 37°C and the clearing reagent was refreshed after 48 h. The clearing procedure was stopped with washes in 0.01% (w/v) sodium azide in PBS. Refractive index matching was achieved by incubating the sections in 25% (w/w) sucrose, 12.5% (w/w) urea, 5% (w/w) triethanolamine and 0.05% (v/v) TritonX-100 for 24 h at 37°C and then in 50% (w/w) sucrose, 25% (w/w) urea, 10% (w/w) triethanolamine and 0.1% (v/v) TritonX-100 for 48 h with refreshing the solution every 24 h.

### Fluorescence *in situ* hybridization (FISH)

40 μm-thick vibratome coronal brain sections were post-fixed in 4% PFA for 15 min and then acetylated in a hydrochloric acid, triethylamine, acetic anhydride solution for 1 min in the dark. Following prehybridization in hybridization buffer (50% formamide, 5x saline-sodium citrate (SSC), 500 μg/ml yeast t-RNA, 1x Denhardt’s solution) at 60°C for 1 h, samples were hybridized with 42 Nm of miR-132 or scramble probe () at 70°C overnight. After sequential washes in 0.2x, 0.5x, 2x SSC buffer and brief incubation in 3% H_2_O_2_, blocking was performed in 0.5% Blocking Reagent (blocking solution) (Roche Diagnostics, Belgium) for 1 h at room temperature. Finally, sections were probed with an antifluorescein HRP-conjugated antibody (Roche Diagnostics, Belgium) in blocking solution for 1 h and signals were developed in TSA Plus Fluorescein reagent (Perkin Elmer, USA). For subsequent immunofluorescent labeling, the sections were processed as described earlier.

### Image acquisition, processing and analysis

For the PCNA staining in human samples, one hippocampal section per case was screened using a widefield Zeiss Axioplan 2 (Zeiss, Belgium). PCNA^+^-nuclei at the subgranular layer of the dentate gyrus within an area that included in total 1000 hematoxylin-stained nuclei were encountered for quantification. Cells with cytoplasmic PCNA^+^-signal were excluded from the analysis, as these may include IBA^+^-proliferating microglia (see Figure S1). The average of the cell counts in the non demented control cohort was used for the normalization of all the samples.

For immunolabelled mouse brain sections and cultured cells, images (z-stacks) were acquired using a Nikon A1R Eclipse Ti confocal microscope (Nikon, Belgium). The FIJI software (Rueden et al., 2017) was employed for image processing and quantification. For BrdU immunostaining in cryosections, three sections per sample were used with 200 μm distance between each two sections. The number of BrdU^+^-cells in the subgranular zone of the dentate gyrus was normalized to the total length of the dentate for each section (BrdU^+^-cells divided by dentate gyrus length). Average values for each sample were calculated by averaging the normalized values of the three sections per sample. For Nestin:GFP, Ki-67 and DCX immunostaining in vibratome-prepared sections, 9-12 sections (with 40 μm distance between each two sections) per sample were used for quantification. Ki-67^+^- (in all the samples) and DCX^+^- (only in the *APP/PS1* and *App^NL-G-F^* dentate gyrus) cells were manually counted per section and average values per sample were calculated by averaging the values of all sections of the same sample. In Nestin:GFP- and DCX- (only in the Nestin:GFP dentate gyrus) immunostained samples, due to the strong signal, the mean intensity values instead of the cell counts were used. Similarly to before, 9-12 sections (with 40 μm distance between each two sections) per sample were used for quantification. Average values per sample were calculated by averaging the values of all sections of the same sample and the values of the control-injected sedentary animals were used for the normalization of all the samples.

For image acquisition of the cleared brain sections, a CSU-X1 spinning disk confocal microscope (tile-scan/z-stack, 10x plan fluor NA 0.3, 488nm laser line) (Nikon, Belgium) was used and 3D rendering of z-stacks was carried using Imaris x64 software (Bitplane, Switzerland). Briefly, GFP^+^-cells were automatically identified as 10 μm ‘spots’ based on size and intensity thresholds were evenly applied for all samples. Reconstructed surfaces were volume-filtered to exclude those not belonging to the dentate gyrus. Spots up to 10 μm close to the dentate gyrus surface were considered for quantification. Values were normalized against the average of the scramble-injected group.

Video files were prepared in Imaris at a 360° angle and at 200 frames of horizontal movement, and saved at 24.000 frames per second and at 640 × 480 VGA.

For detection and quantification of colocalizing pre/post-synaptic puncta, Imaris x64 was used. For assessing dendritic arborization, images of mCHERRY^+^-cells were acquired by regular confocal microscopy. Branching points were counted manually, while the FIJI software was used for quantification of total dendritic length in the same cell. For assessing spine density, dendrite images were acquired using the Zeiss LSM880 Airyscan super-resolution system (Carl Zeiss, Belgium). More specifically, images of mCHERRY-labeled dendritic processes at the outer molecular layer were acquired at 0.2 μm intervals with a 63x oil lens and a digital zoom of 2. Images were subjected to deconvolution and maximum intensity projections of z-series were created with the Imaris x64 software. The length of each dendritic segment was determined by tracing the center of the dendritic shaft and the number of spines was counted automatically from the two-dimensional projections. The linear spine density was calculated by dividing the total number of spines by the length of the dendritic segment.

### RenCell cultures and transfections

RenCell VM human NPCs (Millipore, MA, USA) were cultured according to the manufacturer’s instructions. Differentiation was induced by withdrawing growth factors (FGF-2 and EGF) from culture media. For cell transfections, 40-60% confluent cultures were transfected with miR-132 mimic and control oligonucleotides (at 0.1 pmol or 0.01pmol, depending on the experimental conditions, Dharmacon, Horizon Discovery, UK) or miR-132 antisense inhibitor and control oligonucleotides (at 50 pmol, Dharmacon, Horizon Discovery, UK) using lipofectamine RNAiMAX (ThermoFisher Scientific, Belgium) according to the manufacturer’s instructions. Cells were collected for RNA extraction or fixed for immunostaining three days post transfection. For miR-132 inhibition experiments, differentiation was induced 24 h post transfection.

### Neural induction of human embryonic pluripotent stem cells

Routine culturing and maintenance of human embryonic stem cells (EPSCs) (H9-GFP) (Espuny-Camacho et al., 2013, 2017) was performed using E8-flex growth medium (ThermoFisher Scientific, Belgium). Cells were maintained in a humidified chamber at 37°C with 5% CO_2_ and passaged every 4-5 days with 0.5 mM EDTA. Neural induction of human pluripotent stem cells (NPCs) was carried out according to a previously published protocol (Shi et al., 2012) with some modifications. Briefly, on DIV (days *in vitro*) −2 of neural induction, cells were enzymatically dissociated with StemPro Accutase cell dissociation reagent (Biolegend, CA, USA) and single cells were plated (1.5 × 10^6^/well) in Matrigel-coated (Corning, Sigma-Aldrich, Belgium) 6-well plates in mTeSR1 growth medium (STEMCELL Technologies, France) supplemented with 10 mM Y-27632 ROCK inhibitor (Calbiochem, CA, USA). On DIV0, for initiation of neural induction, the culture medium was replaced with neural maintenance medium supplemented with 1 μM StemMACS LDN-193189 (Miltenyi, Biotec, Netherlands) and 10 μM SB431542 (STEMCELL Technologies, France). Neural induction was carried out for 12 days with daily replacement of growth media. After the neural induction, neuroepithelium was passaged three times using Dispase-II (Roche, Belgium) to enrich for neural rosettes. Culture medium was supplemented with 20 ng/ml FGF (PeproTech, Belgium) during the first Dispase-II step for expansion. Around DIV29, neural rosettes were dissociated using the StemPro Accutase cell dissociation reagent and plated on laminin-coated culture dishes in neural maintenance medium for further maturation. RNA samples were collected on day DIV-2 (PSCs), day DIV29 (NPCs) and DIV60 (neurons).

### Preparation of Aβ_1-42_ oligomers and neuronal precursor cell treatment

Oligomerization and characterization of Aβ_1-42_ was carried out as previously described (Brkic et al., 2015; Kuperstein et al., 2010). Briefly, Aβ_1–42_ (A-1163-1, rPeptide, GA, USA) or scrambled Aβ_1–42_ (A-1004-1, rPeptide, GA, USA) were dissolved at a concentration of 1 mg ml^−1^ in hexafluoroisopropanol (HFIP, Sigma-Aldrich, Belgium). A gentle stream of nitrogen gas was used to evaporate HFIP. The resulting pellet was resuspended in DMSO, and buffer was exchanged with Tris-EDTA using a 5 ml HiTrap desalting column (GE Healthcare, Belgium) according to the manufacturer’s instructions. Determination of the eluted peptide concentration was performed using the Bradford method (Bio-Rad, Belgium). The eluted peptide was allowed to aggregate for 2 h at room temperature. EPSC-derived human neuronal precursor cells were incubated in neural maintenance medium containing 5 μM Aβ_1-42_ or scramble control peptide for 72 h. Six biological replicates per treatment were used.

### Treatment of human neuronal precursor cells with AD and control serum

Serum incubation of human neuronal precursors was performed as previously described (Maruszak et al, 2017, BioRxiv 10.1101/175604). More specifically, the multipotent human hippocampal progenitor/stem cell line HPC0A07/03C (ReNeuron, UK), derived from the first trimester female fetal hippocampal tissue following medical termination and in accordance with UK and USA ethical and legal guidelines, was obtained from Advanced Bioscience Resources (Alameda, CA, USA) and conditionally immortalized by introducing the c-myc-ERTAM transgene, which enables cells to proliferate indefinitely in the presence of epidermal growth factor (EGF), basic fibroblast growth factor (bFGF) and 4-hydroxy-tamoxifen (4-OHT). Removal of these factors induces spontaneous differentiation into neurons, astrocytes or oligodendrocytes. 24 h after seeding, proliferation medium containing EGF, bFGF and 4-OHT was replaced with proliferation medium additionally supplemented with 1% serum. Serum information can be found in Table S4. After 48 h of proliferation, differentiation was induced by replacing culture medium with medium supplemented with 1% serum but devoid of EGF, bFGF and 4-OHT. Each serum treatment was performed in triplicates. Samples were collected for RNA isolation after 7 days of differentiation.

### RNA isolation, reverse transcription and real-time PCR

The dentate gyrus from adult mouse brain was microdissected as previously described (Hagihara et al., 2009). RNA extraction from whole dentate gyri was performed using the miRVana Paris Kit (Life Technologies, Belgium) according to the manufacturer’s instructions. Briefly, tissue was homogenized in 300 μl cell disruption buffer supplemented with protease and phosphatase inhibitors. Following denaturation, addition of acid phenol:chloroform, incubation, and centrifugation, 1.25 volumes of ethanol 100% were added to the aqueous phase. The samples were then loaded on miRVana spin columns and processed according to the manufacturer’s instructions. For isolating RNA from cultured cells, cells were first collected in 1 ml Trizol (Thermo Fisher Scientific, Belgium). Following incubation in chloroform and centrifugation 1.25 volumes of ethanol 100% were added to the aqueous phase and the samples were loaded and processed on miRVana spin columns like before. FACsorted cells were initially collected in RNAprotect Cell reagent (Qiagen, Netherlands) and following brief centrifugation, cell pellets were processed using the miRNeasy Micro kit (Qiagen, Netherlands) according to the manufacturer’s instructions. Reverse transcription of 200 ng (mRNA) or 100 ng (miRNA) RNA was performed using the Superscript II reverse transcriptase (ThermoFisher Scientific, Belgium) for protein-coding transcripts and the Universal cDNA synthesis kit (Exiqon, Qiagen, Denmark) for miRNAs. Real-time semi-quantitative PCR was performed using the SensiFast Sybr No-Rox kit (Bioline, UK) for coding transcripts and the Sybr Green mastermix and LNA PCR primers (Exiqon, Qiagen, Denmark) for miRNAs. Mean expression of two housekeeping genes was used for all normalizations (U6 snRNA, RNU5G for miRNAs, *Actb* and *Gapdh* for murine mRNAs, and *18S* and *GAPDH* for human mRNAs). The primer sequences can be found in Table S3. Cp (crossing points) were determined by using the second derivative method. Fold changes were calculated with the ΔΔ*C*t method (Livak and Schmittgen, 2001).

### Adult dentate gyrus dissociation for Fluorescence Activated Cell Sorting (FACS) of Nestin:GFP^+^ cells

Dentate gyri (each sample consisted of 1 full dentate gyrus; right and left hemisphere) were first dissected and minced in Hibernate A Low Fluorescence (BrainBits, IL, USA) on ice. Tissue dissection was carried out in 20U/ml papain and 50U/ml DNaseI (Worthington, NJ, USA) for 20 min at 37°C. The enzymatic digestion was stopped by incubating the samples in 10 mg/ml ovomucoid solution (Worthington, NJ, USA), cells were dissociated by trituration, filtered through a 70 μm nylon mesh, span down and eventually resuspended in ice-cold Hibernate A. Debris and myelin removal was performed using the adult brain dissociation kit (Miltenyi Biotec, Netherlands) according to the manufacturer’s instructions. Finally, cells were resuspended in ice-cold Hibernate A and dead cells were stained with propidium iodide (ThermoFisher Scientific, Belgium). GFP^+^- and GFP^−^-cells were sorted in a FACS Aria III (BD Biosciences, CA, USA) in RNase-free eppendorfs (in bulk) or in 96-well plates (in single) for downstream analysis.

### Behavioral testing

For contextual fear conditioning assessment, a passive avoidance protocol was employed using a two (an illuminated and a dark one, separated by a guillotine door)-compartment box with a shock grid (Callaerts-Vegh et al., 2015; D’Hooge, 2005). Briefly, dark-adapted, single-housed 9-month old male mice were placed in the illuminated box and latency to enter the dark compartment was recorded. Upon entry into the dark compartment (all four paws inside), the door was closed and a 2 s foot shock (0.2 mA) was applied. The mice were then immediately removed from the box and placed back in their home cage. Twenty four hours later, the dark-adapted mice were again placed in the illuminated box and latency to enter the dark compartment was measured as retention of contextual memory. A maximum of 300 s was noted in case the animal would not enter within the first five minutes.

Pattern separation in 9-month old male mice was measured using an adapted fear conditioning protocol (van Boxelaere et al., 2017; Nakashiba et al., 2012). Briefly, animals were fear conditioned in a specific context (A) by placing them in a conditioning cage (25 × 25 × 25 cm) fitted with stainless steel grid floors to deliver mild foot shocks and located in sound-attenuating cubicles (Panlab Startle & Fear Combined System (Panlab, Spain). Animal movement was monitored by a motion-sensitive floor (the degree of motion could range from 0 to 100) connected to an interfaced computer using Panlab Freezing v1.2.0 software. Freezing was counted if registered movement remained below the arbitrarily defined threshold of 2.5 for at least 1 s. Before testing, animals were placed in a holding area for 30 min for habituation. Contextual fear conditioning consisted of 3 days conditioning in context A, followed by 2 days context testing (A, B, and C, see Figure 7B). In contextual fear acquisition (Days 1-3), mice were placed in context A and after 3 min exploration, a foot shock (2 s; 0.5 mA) was delivered. One minute later, mice were removed from the testing box and placed in their housing cage. Freezing was measured during the 3 min interval preceding the shock.

To determine the specificity of contextual fear conditioning, freezing behavior in contexts A, B and C was recorded during the generalization testing (Days 4–5). On day 4, mice were placed in context A without shocks for 3 min, then removed and placed in a housing cage. 120–150 min later, mice were placed in context B (same features as A, except for an inserted A-frame roof made of black cardboard) and, eventually following the same procedure as before, in context C (change in tactile, olfactory and visual dimension). The testing order between the four groups was randomized to minimize testing time effect, but all animals were tested first in one context before the next context was presented. On day 5, the same procedure was followed but the order of context presentation was changed to B → A →C. Freezing was measured during a 3 min interval and freezing behavior in a similar (B) or a different (C) context was recorded as a measure of discrimination learning (pattern separation).

### Temozolomide (TMZ) treatment

TMZ (Sigma-Aldrich, Belgium) solution (in 10% DMSO) was diluted in 0.9% NaCl to a concentration of 2.5 mg/ml. Control vehicle solution was similarly prepared using 10% DMSO in 0.9% NaCl without TMZ. TMZ or control vehicle were administered i.p. at a dose of 12.5 mg/kg of body weight once a day for 3 consecutive days, followed by 4 days of no injections (one cycle). ICV injections of miR-132 or control mimic were every time performed 2 days after the last TMZ or control vehicle injection of each cycle. After 4 cycles, mice were subjected to the avoidance test.

### Smart-seq2 processing and single-cell library preparation

For FACS isolation of single cells, four miR-132-injected and four control-injected Nestin:GFP male mice were used and 94 cells were initially sorted per mouse. FACS-sorted cells were processed using the Smart-seq2 protocol (Picelli et al., 2013). Briefly, RNA was reverse-transcribed using biotinylated oligo-dT primers (IDT DNA, Belgium) and biotinylated template-switching (TSO) oligonucleotides (Exiqon, Qiagen, Denmark) and subsequently cDNA was preamplified using the KAPA HiFi Hot Start DNA Polymerase (KAPA Biosystems, Roche Diagnostics, Belgium) and biotinylated ISPCR primers (IDT DNA, Belgium). cDNA was purified using AMPURE XP Agencourt beads (Beckman Coulter, France), concentration was calculated with the Quantifluor dsDNA kit (Promega, Netherlands) and adjusted on an Echo 525 Liquid Handler (Labcyte, CA, USA). 100 pg cDNA per cell were used for library preparation with Nextera XT DNA Library Preparation Kit (Illumina, CA, USA). Finally, following purification on magnetic beads the libraries were multiplexed and sequenced using the Illumina NextSeq500 platform (Illumina, CA, USA).

### Data processing

To confirm quality of the raw reads, a FastQC analysis (Version 0.11.5, https://www.bioinformatics.babraham.ac.uk/projects/fastqc/) was performed. Reads were aligned to the mouse mm10 genome (Ensembl Version 88) using STAR (Version 2.5.2 (Dobin et al., 2013)) with default options. A count table was generated from the alignments using Subread/FeatureCounts 1.5.1 (Liao et al., 2014).

The count matrix was analyzed with Seurat 2.3.1 (Butler et al., 2018), using the following workflow: Cells with less than 200 genes or more than 6,000 genes (likely douplets) were removed, as well as cells whose mitochondrial gene expression accounted for more than 20% of the total gene expression of that cell. Genes found in less than 0.5% of cells were also excluded. Following this filtering, 709 of the original 752 cells remained (with a median read depth of 630,131 reads per cell and median of 2,646 genes per cell). Data was log-normalized and scaled by a factor of 10,000. The normalized count data was subsequently regressed on the number of reads. Principal component analysis (PCA) was performed using the top 1,438 highly variable genes. The first 30 PCAs were used to identify clusters (using default parameters). Data was visualized using t-Distributed Stochastic Neighbor Embedding (t-SNE). Cluster markers were determined by a differential expression analysis of each cluster versus all other clusters combined (Table S5; ClusterMarkers). Seurat’s “AddModuleScore” function, with default parameters, was used to calculate gene set scores of cell type-specific markers extracted from previous studies (Table S5; LiteratureMarkers) (Artegiani et al., 2017; Hochgerner et al., 2018; Zeisel et al., 2018). Boxplots of these scores were used to assign each cluster to a cell type (for summary data see Table S5; ModuleScore). We confirmed these assignments by comparing our cluster markers to unique markers of cell type defined by the aforementioned studies studies to ensure that there was no ambiguity in the assignment of cell types. Gene differential expression between experimental (miR-132 overexpression) and control conditions (within each cell population) was performed (Table S6). Differential expression (for analysis of cluster markers and for analysis between controls and experimental conditions) was performed as follows: p-values were calculated using a Wilcoxon rank-sum test (using Seurat’s “FindMarkers” function), and corrected for multiple testing using a Bonferroni correction. Log fold changes were calculated using the natural logarithm. Genes with adjusted p-values of < 0.05 were considered significantly differentially expressed. All sequencing data reported in this study will be deposited in the NCBI Gene Expression Omnibus upon publication.

Functional enrichment analysis was performed by using Ingenuity Pathway Analysis (IPA, Ingenuity Systems Inc., Redwood City, CA, USA). By convention, the 150 transcripts with the most negative (‘Downregulated Transcripts’, left side of x-axis) or most positive (‘Upregulated Transcripts’, right side of x-axis) log fold changes after differential gene expression (comparing miR-132-overexpression to control samples) were selected per cluster for analysis.

