## Supplementary Information for "Impaired adult hippocampal neurogenesis in Alzheimer’s disease is mediated by microRNA-132 deficiency and can be restored by microRNA-132 replacement"

#### Figures

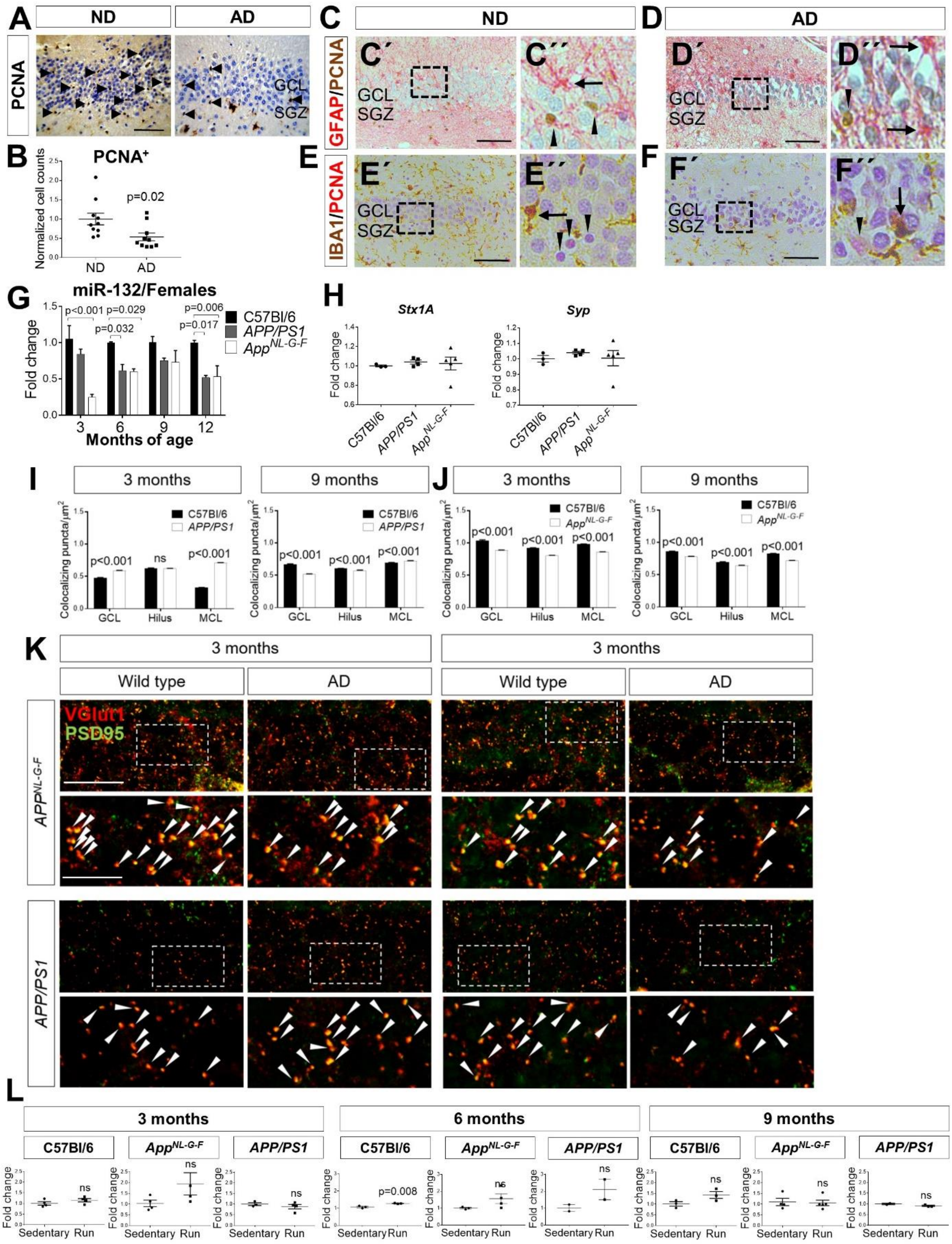

**Figure S1. Related to Figure 1.**

A. Immunohistochemical labeling using the PCNA cell proliferation marker in the dentate gyrus of non-demented (ND) control individuals and Alzheimer's (AD) patients. Arrowheads indicate PCNA$^{+}$-cells at the subgranular zone (SGZ) and the granular cell layer (GCL) as previously reported (Boldrini et al., 2018). Scale bar, 50 µm. B. Quantification of PCNA$^{+}$-cells counts. Values were normalized to 1000 total cell counts per sample (see STAR Methods). N=10 samples per group. C, D. GFAP/PCNA double immunolabeling in the dentate gyrus of non-demented (ND) control individuals (C) and Alzheimer's (AD) patients (D). C'' and D'' show magnified views indicated by dashed rectangles in C' and D', respectively. Arrows indicate GFAP$^{+}$-astrocytes, while arrowheads indicate PCNA$^{+}$-cells. SGZ, subgranular zone; GCL, granular cell layer. E, F. IBA1/PCNA double immunolabeling in the dentate gyrus of ND (E) and AD samples (F). E'' and F'' show magnified views indicated by dashed rectangles in E' and F', respectively. Arrows indicate IBA1$^{+}$-microglia and arrowheads indicate PCNA$^{+}$-cells. Scale bars, 50 µm. G. Semi-quantitative real-time PCR of miR-132 levels in the dentate gyrus of C57Bl/6, *APP/PS1* and *App$^{NL-G-F}$* female mice at 3, 6, 9 and 12 months of age. N=3-6 mice per time point. H. Semi-quantitative real-time PCR of syntaxin 1A (*Stx1A*) and synaptophysin (*Syp*) in the dentate gyrus of 12-month-old C57Bl/6, *APP/PS1* and *App$^{NL-G-F}$* mice. N=4-5 mice per group. I-K. Colocalization of VGlut1$^{+}$-presynaptic and PSD95$^{+}$-postsynaptic puncta in granular cell layer (GCL), hilus and molecular cell layer (MCL) of the dentate gyrus in C57Bl/6, APP/PS1 and *App$^{NL-G-F}$* mice at 3 and 9 months of age. N=3 mice per time point. Arrowheads indicate double-positive puncta. Scale bars, 10 µm (lower magnification); 4 µm (higher magnification). L. Semi-quantitative real-time PCR of miR-212 levels in the dentate gyrus of C57Bl/6, *APP/PS1* and *App$^{NL-G-F}$* sedentary control or running mice at 3, 6 and 9 months of age. N=3-6 mice per group. Values are presented as mean ± SEM. In (G, I, J), two-way ANOVA with Tukey's *post hoc* test for multiple testing were applied, in (H), one-way ANOVA with Tukey's multiple testing, and in (B, L), Student's t-test was used.

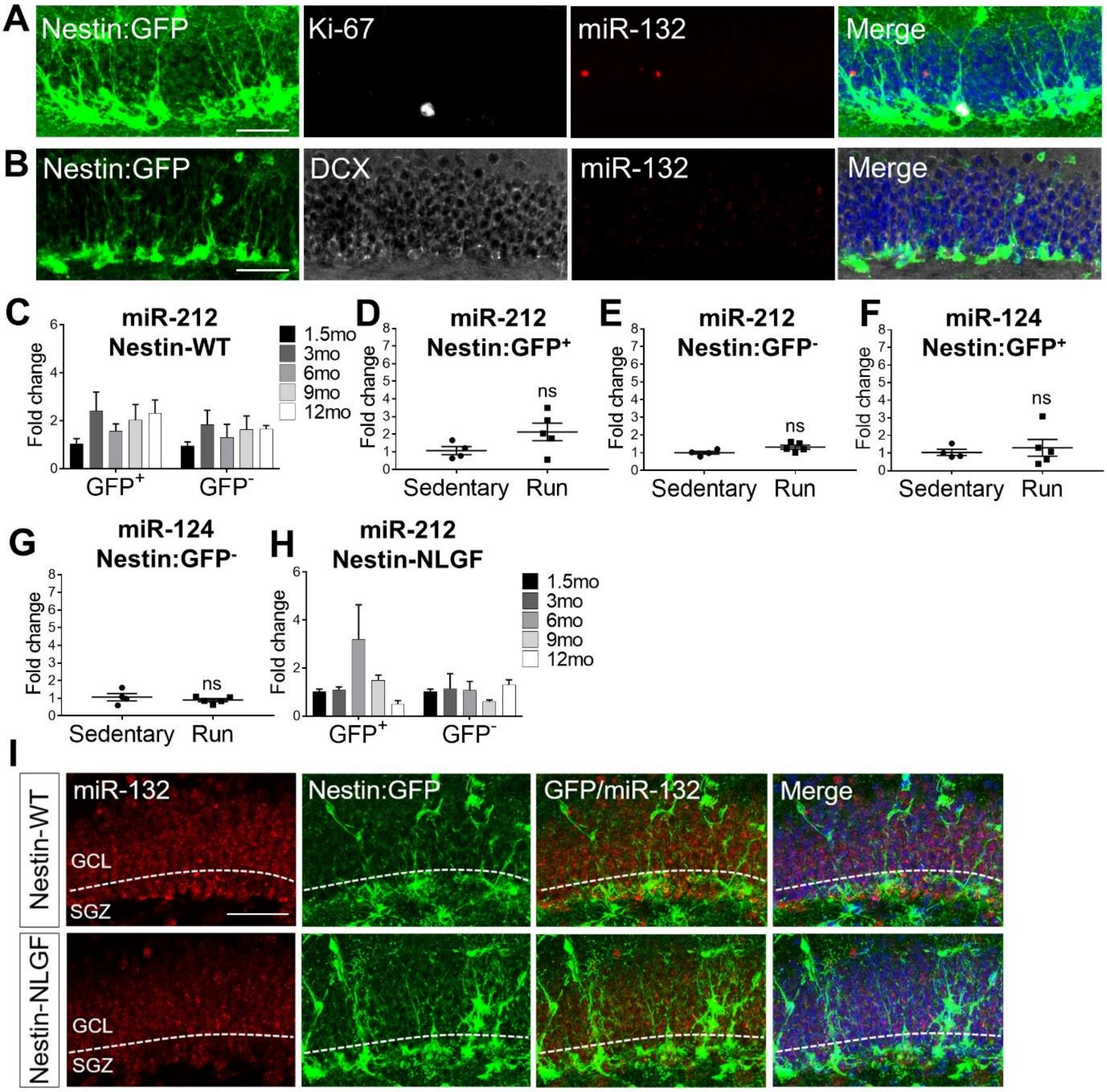

**Figure S2. Related to Figure 2.** A. B. *In situ* hybridization using a scramble control probe combined with Nestin and Ki-67 (A) or Nestin and doublecortin (DCX) (B) immunolabeling in the dentate gyrus of 3-month-old Nestin:GFP mice. Scale bar, 50 µm. C. Semi-quantitative real-time PCR of miR-212 levels in $GFP^+$ or $GFP^-$ populations sorted from the dentate gyrus of Nestin:GFP mice at 1.5, 3, 6, 9 and 12 months of age. N=6-8 mice per time point. D, E. Semi-quantitative real-time PCR of miR-212 levels in $GFP^+$ (D) or $GFP^-$ (E) populations sorted from the dentate gyrus of sedentary control or running Nestin:GFP mice at 3 months of age. N=5-6 mice per group. F, G. Semi-quantitative real-time PCR of miR-124 levels in $GFP^+$ (F) or $GFP^-$ (G) populations sorted from the dentate gyrus of sedentary control or running

Nestin:GFP mice at 3 months of age. N=5-6 mice per group. H. Semi-quantitative real-time PCR of miR-212 levels in $GFP^+$ or $GFP^-$ populations sorted from the dentate gyrus of Nestin-NLGF mice at 1.5, 3, 6, 9 and 12 months of age. N=3-5 mice per time point. I. In situ hybridization for miR-132 in 9-month old Nestin:GFP wild-type (Nestin-WT) and AD (Nestin-NLGF) mice. GCL, granular cell layer; SGZ, subgranular zone. Scale bar, 50 µm. Values are presented as mean ± SEM. In (C, H), two-way ANOVA with Tukey's *post hoc* test for multiple testing were used, while in (D-G), Student's t-test was applied.

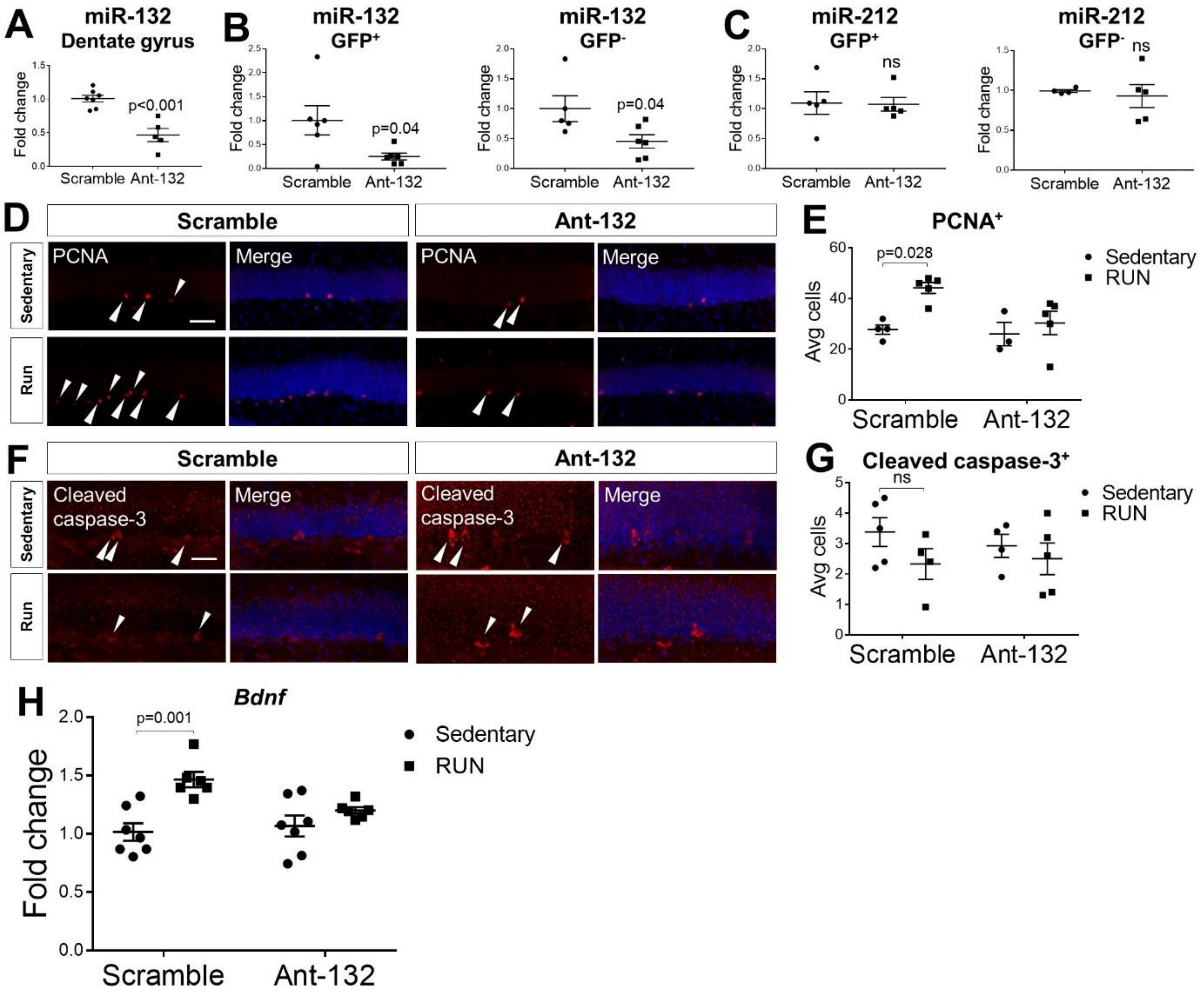

**Figure S3. Related to Figure 3.** A. Semi-quantitative real-time PCR of miR-132 levels in scramble- and Ant-132-injected 9-month-old C57Bl/6 mice prior to running. N=6 mice per group. B. Semi-quantitative real-time PCR of miR-132 levels in $GFP^+$- and $GFP^-$ -cells sorted from the dentate gyrus of Nestin:GFP mice upon miR-132 knockdown prior to running. N=5 mice per group. C. Semi-quantitative real-time

PCR of miR-212 levels in GFP⁺- and GFP⁻ -cells sorted from the dentate gyrus of Nestin:GFP mice upon miR-132 knockdown prior to running. N=5 mice per group. D. PCNA⁺-cells in the subgranular layer of the dentate gyrus of control- or Ant-132-injected, sedentary or running Nestin:GFP mice at 3 months of age. Scale bars, 50 µm. E. Quantification of PCNA⁺-cells in (D). N=3-5 mice per group. F. Cleaved caspase-3⁺-cells in the subgranular layer of the dentate gyrus of control- or Ant-132-injected, sedentary or running Nestin:GFP mice at 3 months of age. Scale bars, 50 µm. G. Quantification of cleaved caspase-3⁺-cells in (F). N=3-5 mice per group. H. Semi-quantitative real-time PCR assessment of the levels of *Bdnf* in the dentate gyrus of control- or Ant-132-injected, sedentary or running Nestin:GFP mice at 3 months of age. N=5-7 mice per group. Values are presented as mean ± SEM. In (A-C), the Student's t-test was applied, while in (E, G, H), two-way ANOVA with Tukey's *post hoc* test for multiple testing were used.

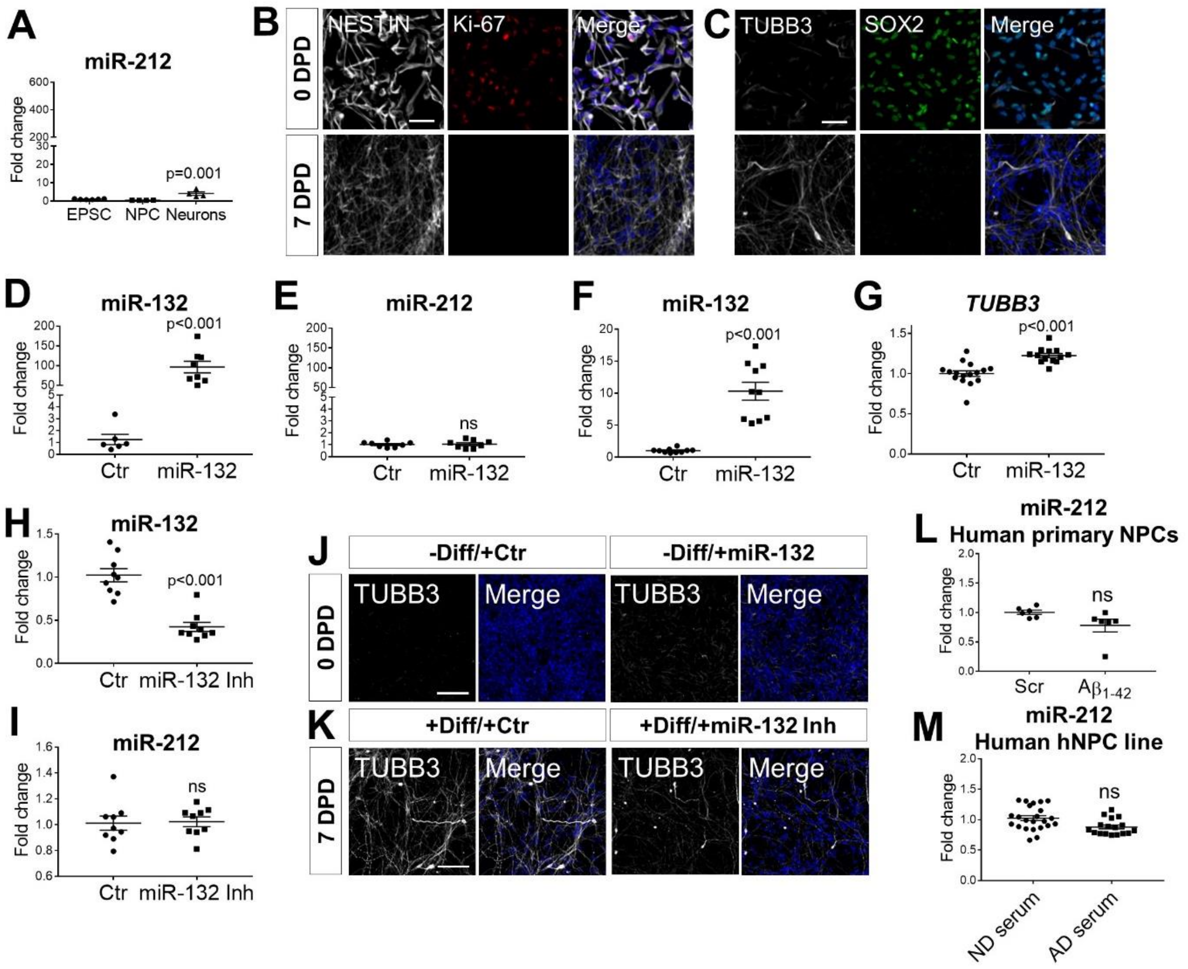

**Figure S4. Related to Figure 4.**

A. Semi-quantitative real-time PCR of miR-212 levels in human EPSCs, NPCs and mature neurons. N=3 independent experiments. B, C. Immunolabeling of NESTIN and Ki-67 (B) or TUBB3 and SOX2 (C) in human neuronal progenitors at 0 and 7 days post differentiation (DPD). Scale bars, 100 µm. D, E. Semi-quantitative real-time PCR of miR-132 (D) and miR-212 (E) levels upon transfection of undifferentiated human neuronal progenitors with 0.1pmol control (Ctr) or miR-132 mimic oligonucleotide (miR-132) 2 days post transfection. N=3 independent experiments. F. Semi-quantitative real-time PCR of miR-132 levels upon transfection of undifferentiated human neuronal progenitors with 0.01pmol control (Ctr) or miR-132 mimic oligonucleotide (miR-132) 2 days post transfection. N=3 independent experiments. G. Semi-quantitative real-time PCR of *TUBB3* levels upon transfection of undifferentiated human neuronal progenitors with 0.01pmol control (Ctr) or miR-132 mimic oligonucleotide (miR-132) 2 days post transfection. N=3 independent experiments. H, I. Semi-quantitative real-time PCR of miR-132 (H) and miR-212 (I) levels upon control or miR-132 inhibitor (miR-132 Inh) transfection of differentiating human neuronal progenitors 2 days post transfection. N=3 independent experiments. J, K. TUBB3 immunostaining in human neuronal progenitors transfected with miR-132 mimic without induction of differentiation (-Diff) (J) or inhibitor with differentiation induction (+Diff) (K) at 0 and 7 days post differentiation (DPD), respectively. Scale bars, 100 µm. L. Semi-quantitative real-time PCR of miR-212 in EPSC-derived human NPCs upon treatment with $A\beta_{1\text{-}42}$ oligomers and a scramble control peptide. N=6 biological replicates. M. Semi-quantitative real-time PCR of miR-212 in an established human hippocampal NPC (hNPC) line (HPC0A07/03CI) upon incubation with AD or control serum. N=23 (ND) and 17 (AD). Values are presented as mean ± SEM. In (A), one-way ANOVA with Tukey's *post hoc* test for multiple testing was employed, while in (D-I, L, M), the Student's t-test was applied.

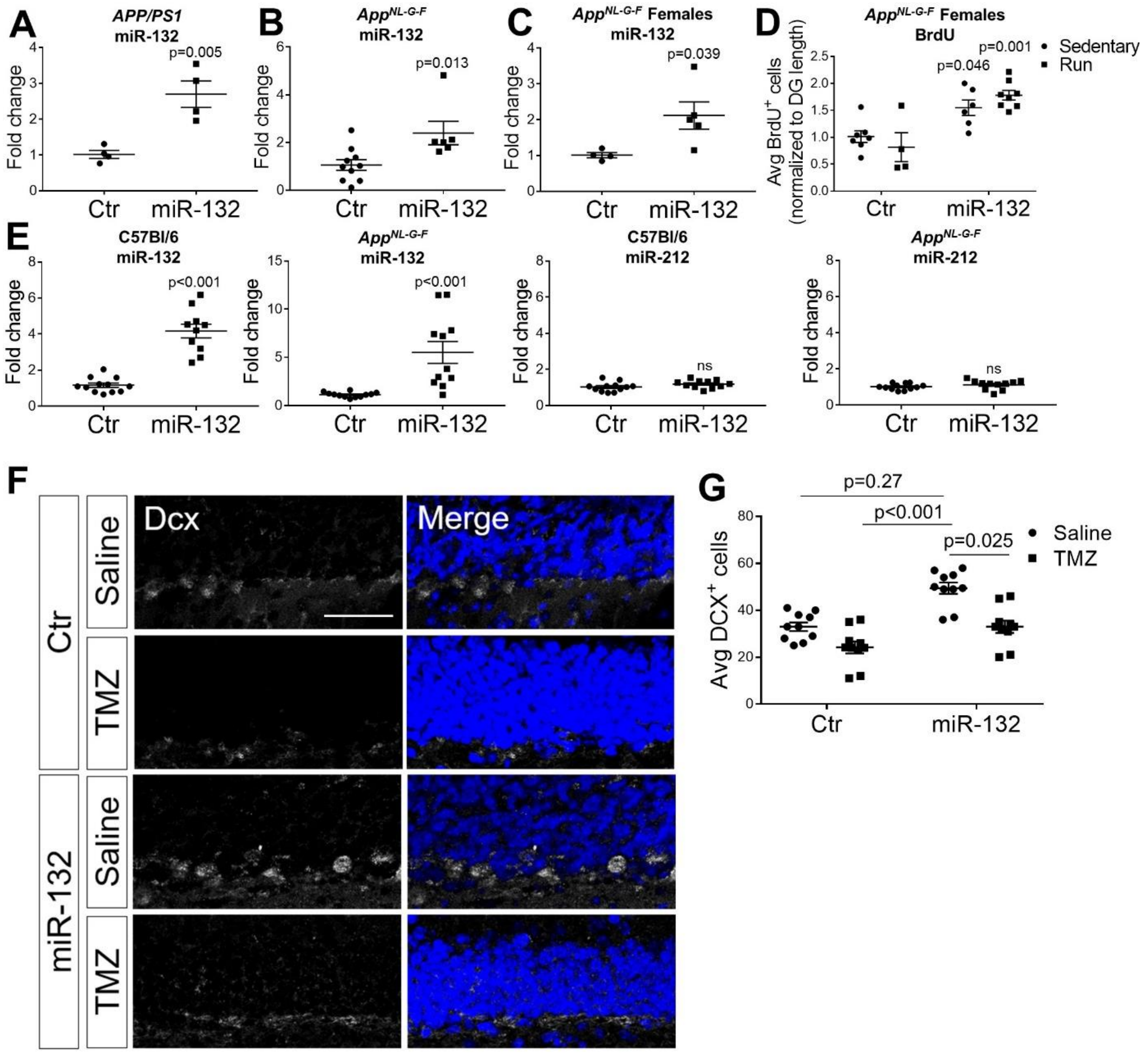

**Figure S5. Related to Figures 5 and 8.** A, B. Semi-quantitative real-time PCR of miR-132 levels in the dentate gyrus of control- or miR-132-injected 9-month-old *APP/PS1* (A) and *App*$^{NL\text{-}G\text{-}F}$ (B) mice prior to running. N=4-10 mice per group. C. Semi-quantitative real-time PCR of miR-132 levels in the dentate gyrus of control- or miR-132-injected *App*$^{NL\text{-}G\text{-}F}$ 9-month-old female mice prior to running. N=4-5 mice per group. D. Quantification of BrdU$^{+}$-cells in the dentate gyrus of control- or miR-132-injected, sedentary or running *App*$^{NL\text{-}G\text{-}F}$ female mice at 9 months of age. N=4-8 mice per group. E. Semi-quantitative real-time PCR of miR-132 and miR-212 levels in the dentate gyrus of C57Bl/6 and *App*$^{NL\text{-}G\text{-}F}$ mice that were exposed to behavioral testing upon miR-132 overexpression. F. Dcx immunolabeling in the dentate gyrus of saline or TMZ-, Ctr- or miR-132-injected *App*$^{NL\text{-}G\text{-}F}$ mice after passive avoidance test completion. G. Quantification of Dcx$^{+}$-cells upon saline or TMZ injections in Ctr- or miR-132-

injected $App^{NL-G-F}$ mice. N=12 mice per group. Scale bar, 50 µm. Values are presented as mean ± SEM. In (A-C, E), the Student's t-test was applied, while in (D, G), two-way ANOVA with Tukey's *post hoc* test for multiple testing were used.

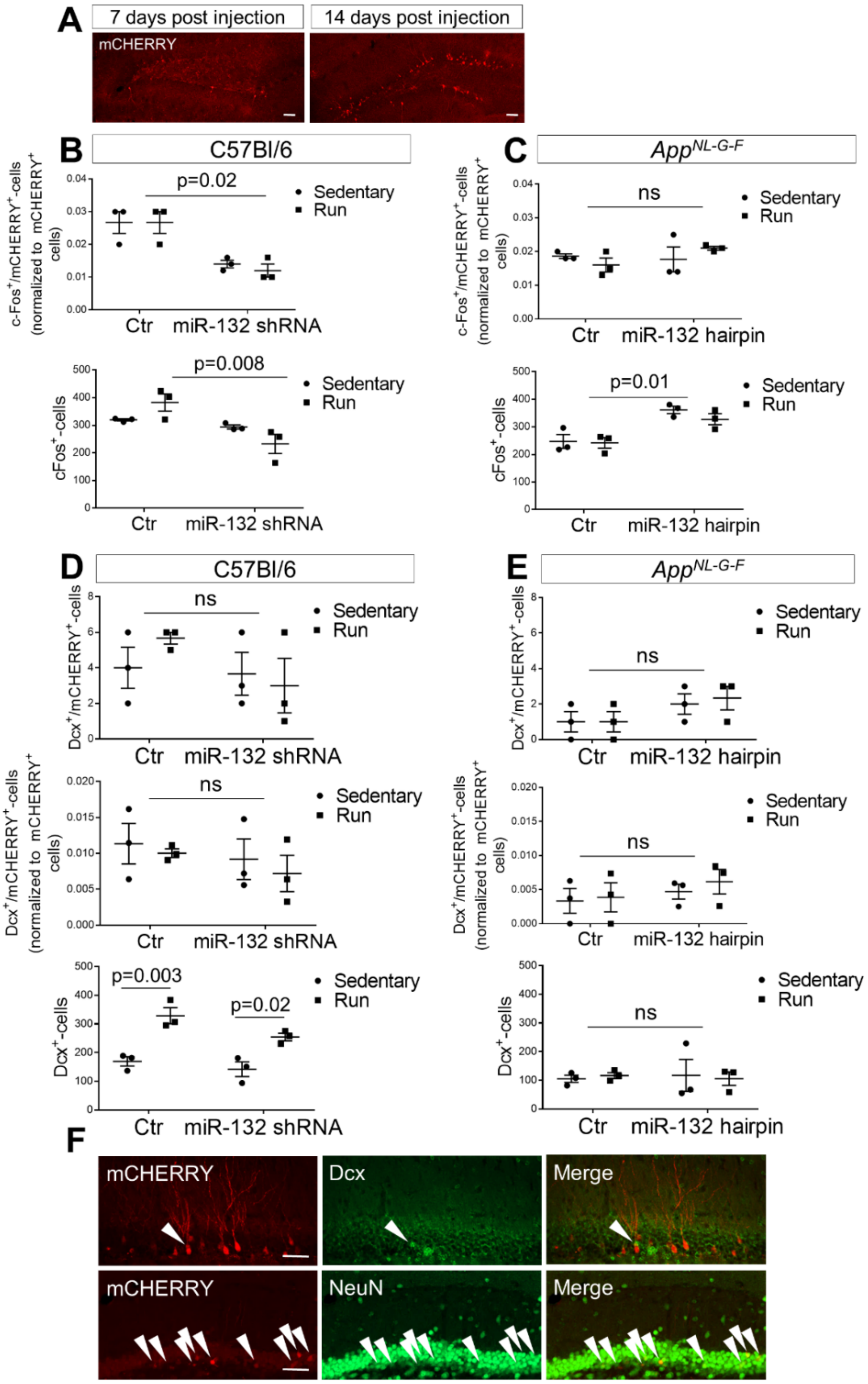

**Figure S6. Related to Figure 6.** A. Representative images of the mCHERRY-labeled 7-day and 14-day old neurons in the dentate gyrus upon injection with the control lentiviral vector. B, C. Quantification of c-Fos$^{+}$/mCHERRY$^{+}$- 4-week old neurons after normalization to the total number of mCHERRY$^{+}$-cells (upper panel) and c-Fos$^{+}$-cells (lower panel) upon miR-132 knockdown in C57Bl/6 mice (B) or miR-132 overexpression in *App*$^{NL\text{-}G\text{-}F}$ animals (C), under sedentary or running conditions. N=3 mice per group. D, E. Quantification of Dcx$^{+}$/mCHERRY$^{+}$ (upper panel), normalized Dcx$^{+}$/mCHERRY$^{+}$ (middle panel) and total Dcx$^{+}$-cells upon miR-132 knockdown in C57Bl/6 mice (D) or miR-132 overexpression in *App*$^{NL\text{-}G\text{-}F}$ animals (E), under sedentary or running conditions. N=3 mice per group. F. Labeling of Dcx$^{+}$/mCHERRY$^{+}$-cells (upper panel) or NeuN$^{+}$/mCHERRY$^{+}$-cells (lower panel) 4 weeks post control lentiviral injections. Arrowheads indicate double-positive cells in each panel. Scale bars, 50 µm. Values are presented as mean ± SEM. In (B-E), two-way ANOVA with Tukey's *post hoc* test for multiple testing were used.

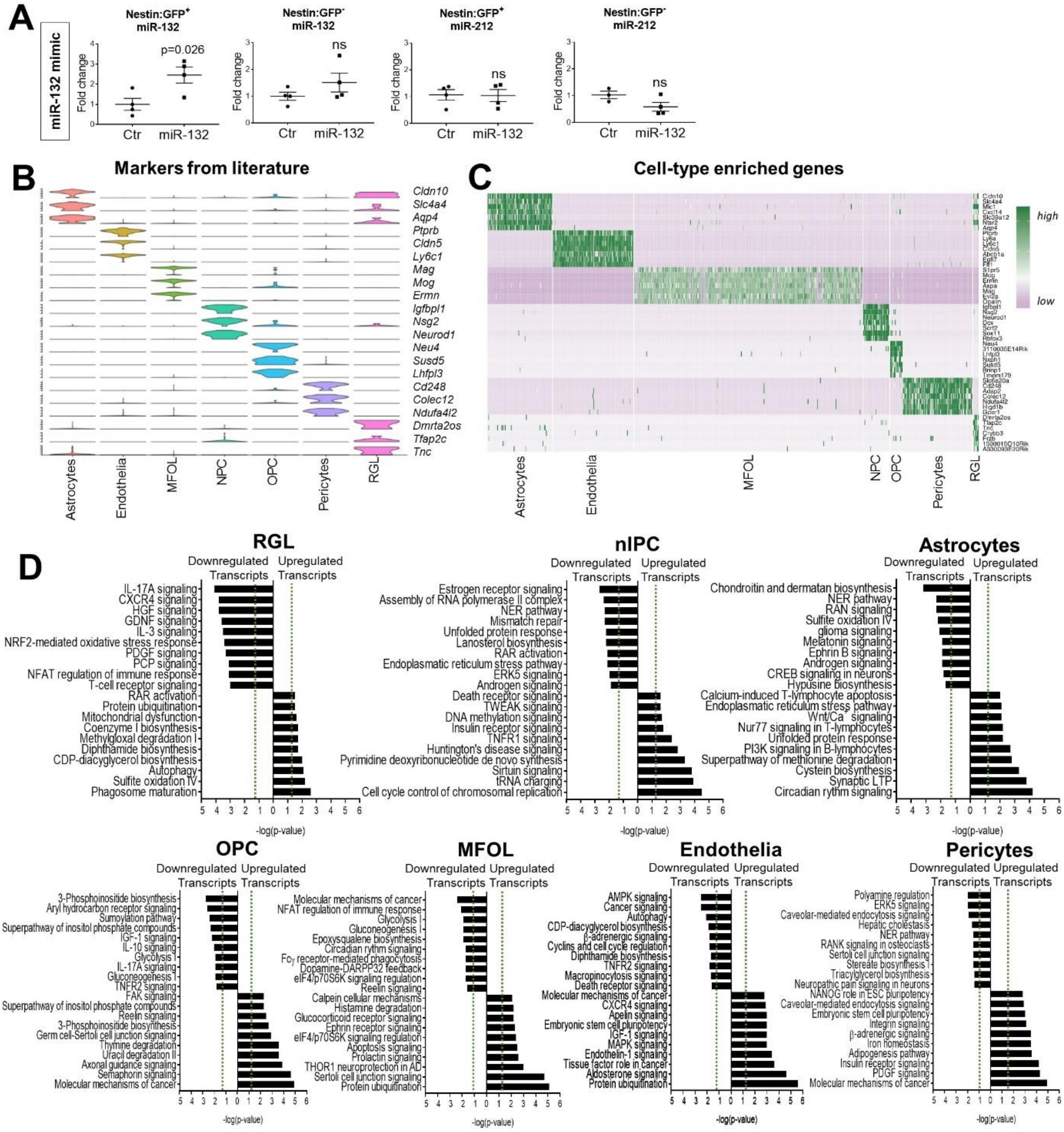

**Figure S7. Related to Figure 7.** A. Semi-quantitative real-time PCR of miR-132 and miR-212 levels in GFP+ or GFP- populations sorted from the dentate gyrus of Nestin:GFP mice at 3 months of age upon control- or miR-132-injection. N=4 mice per group; Student's t-test was used. B. Cell type-specific gene expression of markers derived from previously published datasets. C. Heatmap depicting the expression of the top ten enriched genes per cell type across all identified cells. Each row represents a single cell and each column represents a gene. The expression is normalized by gene. For all significantly upregulated genes per cell cluster, see Table S5. D. Gene ontology enrichment analysis per cluster. Top ten significantly enriched pathways per cluster are indicated for both the 150 most increasing genes (right side of x-axis) and the 150 most decreasing genes (left side of x-axis) upon miR-132 overexpression. The significance values for the canonical pathways were calculated by Fisher's exact test, right-tailed and the threshold for –log(p-value) was set at 1.3 (dashed line).

### Tables

**Table S1. Cases used for immunohistochemical analysis of human dentate gyrus. Related to Figures 1 and S1.** Histopathological and clinical scores were determined as previously published and recommended (Rijal Upadhaya et al., 2014). F, female; M, male; Age in years; PMI, postmortem interval (in hours); Aβ-MTL (medial temporal lobe) phase (Thal et al., 2000); Braak-NFT (neurofibrillary tangle) stage (Braak et al., 2006); CERAD (Consortium to Establish a Registry for Alzheimer's Disease)-plaque score for neuritic plaque density (Mirra et al., 1991); NIA-AA (National Institute on Aging–Alzheimer's Association) degree of Alzheimer's pathology (Hyman et al., 2012); CDR (Clinical Dementia Rating) (Morris, 1993); N.A., not available.

**Table S2. Antibodies used in the study.**

| Name | Host | Dilution | Supplier | Catalog number |
|---|---|---|---|---|
| Anti-BrdU | Rat | 1:100 | Abcam | ab6326 |
| Anti-GFP | Chicken | 1:500 | Abcam | ab13970 |
| Anti-Ki-67 | Rabbit | 1:500 | Genetex | GTX16667 |
| Anti-DCX | Rabbit | 1:250 | Abcam | ab18723 |
| Anti-TUBB3 | Mouse | 1:200 | Biolegend | 801202 |
| Anti-GFAP | Rabbit | 1:500 | DAKO | IS524 |
| Anti-PCNA | Mouse | 1:200 | DAKO | M0879 |

| | | | | |
|---|---|---|---|---|
| Anti-IBA1 | Rabbit | 1:500 | WAKO | 019-19741 |
| Anti-NESTIN | Mouse | 1:200 | R&D Systems | MAB1259 |
| Anti-SOX2 | Goat | 1:200 | Santa Cruz | sc-365823 |
| Anti-Cleaved caspase-3 | Rabbit | 1:400 | Cell Signaling | 9661 |
| Anti-VGlUT1 | Guinea pig | 1:1000 | Millipore | AB5905 |
| Anti-PSD95 | Mouse | 1:200 | Thermo Scientific | 7E3-1B8 |
| Anti-mCHERRY | Rabbit | 1:500 | Clontech | 632496 |
| Anti-mCHERRY | Goat | 1:200 | LifeSpan | LS-C204207 |
| Anti-c-FOS | Rabbit | 1:300 | Millipore | PC38 |
| Anti-NeuN | Mouse | 1:200 | Millipore | MAB1281 |

**Table S3. Primers used in the study.**

| **Primer (microRNAs)** | **Target sequence** |
|---|---|
| mmu-miR-132-3p | UAACAGUCUACAGCCAUGGUCG |
| mmu-miR-212-3p | UAACAGUCUCCAGUCACGGCCA |
| mmu-miR-124 | UAAGGCACGCGGUGAAUGCC |
| **Primer (messenger RNAs)** | **Sequence (5' - 3')** |
| *TUBB3* Forward (Homo sapiens) | CCTCCGTGTAGTGACCCTT |
| *TUBB3* Reverse (Homo sapiens) | GGCCTTTGGACATCTCTTCAG |
| *MKI67* Forward (Homo sapiens) | GAGGTGTGCAGAAAATCCAAA |
| *MKI67* Reverse (Homo sapiens) | CTGTCCCTATGACTTCTGGTTGT |
| *GAPDH* Forward (Homo sapiens) | TCAAGAAGGTGGTGAAGCAGG |
| *GAPDH* Reverse (Homo sapiens) | ACCAGGAAATGAGCTTGACAAA |
| *18S* Forward (Homo sapiens) | TTCGAGGCCCTGTAATTGGA |

| | |
|---|---|
| *18S* Reverse (Homo sapiens) | GCA GCAACTTAATATACGCTAT |
| *DOCK1* Forward (Homo sapiens) | CTGGCCTTGTTTGTGAACCT |
| *DOCK1* Reverse (Homo sapiens) | TCTTTGCTTCCGAGGTCAGT |
| *EPHB3* Forward (Homo sapiens) | GCTGTGATGACAACGTGGAG |
| *EPHB3* Reverse (Homo sapiens) | TAGTGTGGGCACTTCAGACG |
| *BTG2* Forward (Homo sapiens) | AGGAGGCACTCACAGAGCAC |
| *BTG2* Reverse (Homo sapiens) | CGGTAGGACACCTCATAGGG |
| *CAMK1* Forward (Homo sapiens) | CAAAGATTTCATCCGGCACT |
| *CAMK1* Reverse (Homo sapiens) | TTGAAGGCTTGCTTCCACTT |
| *RAC1* Forward (Homo sapiens) | TTTGAAAATGTCCGTGCAAA |
| *RAC1* Reverse (Homo sapiens) | TCGCTTCGTCAAACACTGTC |
| *JUN* Forward (Homo sapiens) | CCCCAAGATCCTGAAACAGA |
| *JUN* Reverse (Homo sapiens) | CCGTTGCTGGACTGGATTAT |
| *CD320* Forward (Homo sapiens) | GGCTGTGGAACCAATGAGAT |
| *CD320* Reverse (Homo sapiens) | CAGAGGAGGATGTGGCATTC |
| *SESN3* Forward (Homo sapiens) | CCCTTGACAAGAGGACCAAG |
| *SESN3* Reverse (Homo sapiens) | CCATGCGCAACATGTAAAAC |
| *MSN* Forward (Homo sapiens) | ATCACTCAGCGCCTGTTCTT |
| *MSN* Reverse (Homo sapiens) | CCCACTGGTCCTTGTTGAGT |
| *TMCC3* Forward (Homo sapiens) | CATGAGACAGCCAACCTGAA |
| *TMCC3* Reverse (Homo sapiens) | AGAATGTGGCAGCGACTCTT |
| *TNIK* Forward (Homo sapiens) | AAATCTTACGGGGGCTGAGT |
| *TNIK* Reverse (Homo sapiens) | TGCCATCCAGTAGGGAGTTC |
| *PDHX* Forward (Homo sapiens) | CCAAAGACGTAGGTCCTCCA |
| *PDHX* Reverse (Homo sapiens) | TGGCTAGCATCCAGTGAGTG |

| | |
|---|---|
| *DENR* Forward (Homo sapiens) | CCAAGGAACAGTGCCAAGTT |
| *DENR* Reverse (Homo sapiens) | GACCCCTTCCACCTCTCTTC |
| *FOS* Forward (Homo sapiens) | AGAATCCGAAGGGAAAGGAA |
| *FOS* Reverse (Homo sapiens) | CTTCTCCTTCAGCAGGTTGG |
| *UBR5* Forward (Homo sapiens) | ATTGCTGTGCCCTTTACACC |
| *UBR5* Reverse (Homo sapiens) | TGAAAATGCAACTGCTCCAG |
| *COA3* Forward (Homo sapiens) | GACTCGGGAGAAGCTGACAC |
| *COA3* Reverse (Homo sapiens) | CTTTGGCCTCGTCTTCTAGC |
| *ZMYM5* Forward (Homo sapiens) | TATCGAATGGGGACTTCCTG |
| *ZMYM5* Reverse (Homo sapiens) | CCCCTGGCTGTTTCTGATTA |
| *MGA* Forward (Homo sapiens) | CTCTCCAAATGTCCCTGGAA |
| *MGA* Reverse (Homo sapiens) | TGTTGTCCCATCAACTGCAT |
| *CORO7* Forward (Homo sapiens) | CTGGTGTACTGGGCATTGTG |
| *CORO7* Reverse (Homo sapiens) | GCAGTCGCCAGAGTTTTACC |
| *GSK3B* Forward (Homo sapiens) | TTCCTTTGGAATCTGCCATC |
| *GSK3B* Reverse (Homo sapiens) | ACAGCTCAGCCAACACACAG |
| *EFHD1* Forward (Homo sapiens) | GGCAAAGCTTTCTGAGATCG |
| *EFHD1* Reverse (Homo sapiens) | GTTGGCCTTGAGTTTCTGGA |
| *ZHX3* Forward (Homo sapiens) | CTGAGCAGCATTCCAACGTA |
| *ZHX3* Reverse (Homo sapiens) | AACCGTAATTGTGGGCTGAG |
| *CDC40* Forward (Homo sapiens) | CCTCAGTGCAGCCTATGACA |
| *CDC40* Reverse (Homo sapiens) | GGTGTTGACAGCTCCCAAAT |
| *PPP2R5C* Forward (Homo sapiens) | AGCCAATCCCCAGTACACAG |
| *PPP2R5C* Reverse (Homo sapiens) | ACGGTCCTTCTTCGGATCTT |
| *AAK1* Forward (Homo sapiens) | GCACCAGAAATGGTCAACCT |

| | |
|---|---|
| *AAK1* Reverse (Homo sapiens) | GGCAGTGCATGTCTTGAGAA |
| *PSME4* Forward (Homo sapiens) | TTCAGCCACAGCTGAATTTG |
| *PSME4* Reverse (Homo sapiens) | ACCATGCGACCTGCTACTCT |
| *SLC1A4* Forward (Homo sapiens) | AGCTCAACGCAGGACAGATT |
| *SLC1A4* Reverse (Homo sapiens) | GCTTCCACTTTCACCTCAGC |
| *UTRN* Forward (Homo sapiens) | AGTCACCATAGACGCCATCC |
| *UTRN* Reverse (Homo sapiens) | GTCCTCAGCAGAAAGCAACC |
| *FAM168A* Forward (Homo sapiens) | TTACCCCACAGCCTATCCAG |
| *FAM168A* Reverse (Homo sapiens) | AGTGCCACAGGAAGACGAGT |
| *CLDN10* Forward (Homo sapiens) | GATCATCGCCTTCATGGTCT |
| *CLDN10* Reverse (Homo sapiens) | GCTGACAGCAGCGATCATAA |
| *CHMP1A* Forward (Homo sapiens) | AAGAAGGCGGAGAAGGACTC |
| *CHMP1A* Reverse (Homo sapiens) | GGTCACCCCCTTCATAGTCA |
| *GCSH* Forward (Homo sapiens) | GAAGCGTTGGGAGATGTTGT |
| *GCSH* Reverse (Homo sapiens) | CCTGGATTTTCTGCAAGAGC |
| *RASA1* Forward (Homo sapiens) | AGCGAAAAACGAGCTACCAA |
| *RASA1* Reverse (Homo sapiens) | CGAAGGCGTTTATTGGATGT |
| *EXOC6B* Forward (Homo sapiens) | GAACTGTGGGAAATGGCACT |
| *EXOC6B* Reverse (Homo sapiens) | ACCTGCCCACTTCTTTAGCA |
| *ORAI3* Forward (Homo sapiens) | GAAGCTGTGAGCAACATCCA |
| *ORAI3* Reverse (Homo sapiens) | AACTTGACCCAACCAACCAG |
| *HDAC3* Forward (Homo sapiens) | TGGCTTCTGCTATGTCAACG |
| *HDAC3* Reverse (Homo sapiens) | GTTGATAACCGGCTGGAAAA |
| *USE1* Forward (Homo sapiens) | GCAGCTGAGCTAGACCTCGT |
| *USE1* Reverse (Homo sapiens) | GTCCGCCATTTTCAGTGAGT |

| | |
|---|---|
| *SLC25A12* Forward (Homo sapiens) | CTGAGGGGGCCTTACCTTAC |
| *SLC25A12* Reverse (Homo sapiens) | CATTCGGGTCTTCACCAGAT |
| *SLC35B4* Forward (Homo sapiens) | ATTCCAGGCATTTGTGTGGT |
| *SLC35B4* Reverse (Homo sapiens) | GATGGATTCCAGGCATTTGT |
| *PLXNB2* Forward (Homo sapiens) | TCCAGCTCCTCCTTAGACGA |
| *PLXNB2* Reverse (Homo sapiens) | AGGTTCTTGCCCTGGAAGTT |
| *ADAMTS1* Forward (Homo sapiens) | ACGAGTGCGCTACAGATCCT |
| *ADAMTS1* Reverse (Homo sapiens) | TCCTTGCACACAGACAGAGG |
| *ATP6V0A1* Forward (Homo sapiens) | GTCACCGACCTTGACTCCAT |
| *ATP6V0A1* Reverse (Homo sapiens) | AATGCCATGACCGAAGTCTC |
| *SIPA1L3* Forward (Homo sapiens) | ACCCCTGCACTGATAACGTC |
| *SIPA1L3* Reverse (Homo sapiens) | GGTGTGGAACTTGTCGGACT |
| *CBFA2T2* Forward (Homo sapiens) | CACGAACACCTTCTGCTCAA |
| *CBFA2T2* Reverse (Homo sapiens) | GTGGAGGGGTAGGATGGAAT |
| *KAT7* Forward (Homo sapiens) | GAAATGCGCCTTCTTCTGAG |
| *KAT7* Reverse (Homo sapiens) | TGTGCTCTCACCTTGCATTC |
| *CIAO1* Forward (Homo sapiens) | CCCACACACAGGATGTCAAG |
| *CIAO1* Reverse (Homo sapiens) | GACGCCAGATACGCACAGTA |
| *GAPVD1* Forward (Homo sapiens) | TGTTAGGCAGTTTGCTGTGC |
| *GAPVD1* Reverse (Homo sapiens) | CCCAGTCCAGAACAGCTCTC |
| *ABCF1* Forward (Homo sapiens) | GACCACACTCCTCAAGCACA |
| *ABCF1* Reverse (Homo sapiens) | CCGCAATTCCTCATACACCT |
| *CRKL* Forward (Homo sapiens) | ATCTGTCTCAGCACCCAACC |
| *CRKL* Reverse (Homo sapiens) | GGCACTCCACCACTGTTCTT |
| *DIP2A* Forward (Homo sapiens) | CCTGTGATGTTCATGGTTGC |

| | |
|---|---|
| *DIP2A* Reverse (Homo sapiens) | TCTGACAAGCGTCTGTGGTC |
| *MDM4* Forward (Homo sapiens) | TGGGTACTGCCATTGTTTCA |
| *MDM4* Reverse (Homo sapiens) | CACTGCCACTCATCCTCAGA |
| *ASCC2* Forward (Homo sapiens) | TCAGGGCTTCATCGAAGAGT |
| *ASCC2* Reverse (Homo sapiens) | CACCGATGGGTCTTTAGCAT |
| *Stx1A* Forward (Mus musculus) | AGCACAACATACCCTGTGG |
| *Stx1A* Reverse (Mus musculus) | GGTCTTCCAGTTACCCGACA |
| *Syp* Forward (Mus musculus) | TGCCAACAAGACGGAG |
| *Syp* Reverse (Mus musculus) | GGCGGATGAGCTAACT |
| *Pax6* Forward (Mus musculus) | AACAACCTGCCTATGCAACC |
| *Pax6* Reverse (Mus musculus) | ACTTGGACGGGAACTGACAC |
| *Mki67* Forward (Mus musculus) | GTCCCTCAAAAGCAGACGAG |
| *Mki67* Reverse (Mus musculus) | GCTGCTGCTTCTCCTTCACT |
| *Dcx* Forward (Mus musculus) | TCCCCAACACCTCAAAAGAC |
| *Dcx* Reverse (Mus musculus) | ATGGAATCGCCAAGTGAATC |
| *Actb* Forward (Mus musculus) | TCCTCCCTGGAGAAGAGCTA |
| *Actb* Reverse (Mus musculus) | GCAATGATCTTGATCTTC |
| *Gapdh* Forward (Mus musculus) | TTGATGGCAACAATCTCCAC |
| *Gapdh* Reverse (Mus musculus) | CGTCCCGTAGACAAAATGGT |

**Table S4. Information on serum samples used in the '*in vitro* parabiosis' assay. Related to Figures 4 and S4.** Blood samples were drawn and biochemically characterized as previously described (Johansson et al., 2013). n, sample size; Age in years; SD, standard deviation; T-tau, total tau; P-tau, tau phosphorylated at threonine 181.

**Table S5. Cell-type mapping. Related to Figures 7 and S7.** Seurat's "AddModuleScore" function, with default parameters, was used to calculate gene set scores of cell type-specific markers extracted from previous studies (LiteratureMarkers). Boxplots of these scores were used to assign each cluster to a cell type (summary data provided under ModuleScore). Cluster markers were determined by differential expression analysis of each cluster versus all other clusters combined (ClusterMarkers). Qu, quartile; RGL, radial glia-like neural stem cells; NPC, neuronal intermediate progenitor cells; OPC, oligodendrocyte precursor cells; MFOL, myelin-forming oligodendrocytes; p_val, p-value; avg_logFC, average of log fold change; p_val_adj, adjusted p-value using the Bonferroni correction.

**Table S6. Differential expression analysis in single Nestin:GFP$^{+}$-niche cells. Related to Figures 7 and S7.** Differential expression analysis was performed between miR-132-overexpression and control group in RGL (radial glia-like neural stem cells), NPC (neuronal intermediate progenitor cells), astrocytes, OPC (oligodendrocyte precursor cells), MFOL (myelin-forming oligodendrocytes), endothelial cells and pericytes. p_val, p-value; avg_logFC, average of log fold change; pct1, percentage of cells in first group (miR-132 overexpression group); pct2, percentage of cells in second group (control group); p_val_adj, adjusted p-value using the Bonferroni correction; DE, differential expression.
